# A spectral partial information decomposition framework for quantifying information about cognitive variables in oscillatory brain networks

**DOI:** 10.64898/2026.05.13.724846

**Authors:** Vinicius Lima, Daniele Marinazzo, Andrea Brovelli

## Abstract

Neural oscillations are thought to play a central role in encoding and transmitting cognitive information across large-scale brain networks, yet the relative contributions of phase synchrony and amplitude co-modulations to distributed coding remain unclear. A key obstacle is the absence of tools that can simultaneously quantify task-relevant information in the frequency domain and disentangle its phase and amplitude co-modulation components across pairwise and higher-order interactions. Here, we introduce a spectral partial information decomposition framework (named NeOPID) for quantifying information about cognitive variables in amplitude co-modulation and phase contributions, and to quantify redundant and synergistic information in brain relations, from pairwise to higher-order interactions. We validated the approach on Kuramoto and Stuart-Landau oscillator networks, including a whole-brain model constrained by macaque anatomical connectivity. NeOPID accurately recovers ground-truth encoding schemes and reveals that phase relations and amplitude co-modulations act as complementary coding channels with both redundant and synergistic components. NeOPID further extends this decomposition to higher-order functional interactions enabling the characterization of how cognitive information is collectively distributed across multiple oscillatory edges via redundant and synergistic encoding. To illustrate biological applicability, we applied NeOPID to local field potentials (LFPs) recorded from the macaque fronto-parietal network during a working memory task. In this dataset, NeOPID identified beta-band amplitude co-modulations as the primary carrier of stimulus information, and revealed that higher-order phase interactions exhibit both redundant and synergistic structure during the memory delay. These results establish NeOPID as a principled tool for dissecting the informational architecture about cognitive processes of oscillatory brain networks.

**Significance statement:** Understanding how the brain coordinates large-scale communication to support cognition remains a fundamental challenge in neuroscience. Characterizing the distinct roles of phase synchrony and amplitude co-modulation has been hindered by a lack of computational approaches capable of separating these components across pairwise and higher-order interactions. Here, we introduce NeOPID (neural spectral Partial Information Decomposition), a framework designed to isolate phase and amplitude co-modulation contributions and quantify redundant and synergistic information contained in pair-wise and higher order functional interactions. NeOPID was validated on simulated oscillator models and on macaque local field potentials (LFPs) during a working memory task. Overall, the results establish NeOPID as a principled tool for dissecting the informational architecture underlying cognitive processes in oscillatory brain networks.

## INTRODUCTION

A central hypothesis in neuroscience posits that cognitive functions emerge from the coordinated activity and interactions of widespread neural populations (Varela et al., 2001b; Bressler and Menon, 2010) resulting in self-organized networks encoding and routing information relevant for cognitive functions (Battaglia and Brovelli, 2020; Deco et al., 2015). An increasing number of studies indicate that neural functional interactions, quantified via statistical relationships between the activity of different brain regions, reflect task-relevant information (Panzeri et al., 2022; Combrisson, 2022), with evidence that this information is structured into redundant and synergistic components (Varley et al., 2023a; Luppi et al., 2024; Combrisson et al., 2024, 2025). Redundancy captures the shared information about cognitive variables in both brain regions, or neuronal populations. In contrast, synergy reflects the additional information present only in the joint observation of both regions, which cannot be obtained from either one individually. Current hypotheses suggest that redundant information may enhance robustness by ensuring that critical information is consistently duplicated and distributed to downstream readout (Valente, 2021; Combrisson et al., 2024). Synergistic information may be associated with flexibility and provide functional advantage, because it would enable the full exploitation of possible combinations of neurons or brain areas, making their informational capacity to scale exponentially with the system size (Rosas et al., 2020). Indeed, synergistic interactions have been reported between distant associative regions during high-level cognition (Luppi et al., 2022; Varley et al., 2023a; Combrisson et al., 2024, 2025) and, at the microscale, in populations of neurons within a cortical column of the visual cortex and across areas of the visuomotor network (Nigam et al., 2019).

Additionally, several studies suggest that task-relevant information is contained in collective and higher-order statistical dependencies across multiple neurons or brain regions, beyond pairwise relations (Martignon, 2000; Yu, 2011; Shahidi et al., 2019; Chelaru, 2021; Panzeri et al., 2022; Varley et al., 2023b; Santoro et al., 2023). Generally, higher-order interactions are at the core of complex systems (Crutchfield, 1994; Battiston et al., 2020; Battiston, 2021) and they are thought to support collective behavior in social systems (Benson et al., 2016), ecology (Grilli et al., 2017; Levine et al., 2017) and biology (Sanchez-Gorostiaga et al., 2019). In brain dynamics, higher-order statistical dependencies among multiple neurons or brain regions may reflect complex coordinated activity associated with emergent functional properties, such as information integration and cognitive flexibility (Schneidman et al., 2006; Kö ster et al., 2014; Chelaru, 2021; Santoro et al., 2024). Indeed, a recent study has shown that an important goal-directed learning signal, such as information gain, is reflected in synergistic and higher-order functional interactions between distant brain areas, and that this information is directionally structured toward key brain structures such as the prefrontal reward circuits (Combrisson et al., 2025). Taken together, the tradeoff between synergistic and redundant information within high-order functional interactions may support an efficient and distributed representation of information relevant to cognitive functions.

A large body of evidence also suggest that neural oscillations play a crucial role in brain-wide interactions and cognitive processes, such as selective attention (Womelsdorf and Fries, 2007; Womelsdorf et al., 2007; Bosman et al., 2012; Bonnefond et al., 2017; van Kerkoerle et al., 2014; Fries et al., 2001) and working memory (Salazar et al., 2012; Miller et al., 2018; Lundqvist et al., 2016; Dotson et al., 2018). Several accounts have been proposed to explain the mechanisms supporting neural interactions, including phase synchronization between population-level activities (Singer and Gray, 1995; Engel et al., 2001; Varela et al., 2001b; Fries, 2015, 2009; Buzsáki and Draguhn, 2004; Siegel et al., 2012), frequency-specific feedforward and feedback influences (Vezoli et al., 2021; Bastos et al., 2015; Xiong et al., 2024), and entrainment via the resonance between the sender and receiver oscillatory transient dynamics (Vinck et al., 2023, 2025). Although these accounts differ in their specific mechanisms, they all highlight the importance of phase relations between neural oscillations and co-modulations in transient dynamics. These notions, and the concept of intrinsic coupling modes (ICMs), have been proposed to capture the dynamics of ongoing neuronal interactions at multiple spatial and temporal scales, via phase coupling for fast synchronization and envelope/amplitude correlation for slow power fluctuations (Engel et al., 2013). Progress toward a unified interpretation is, however, limited by the lack of tools to disentangle the role of phase and amplitude relations in cognition.

The aim of our study was to disentangle spectrally-resolved information in transient amplitude co-modulations and phase relations, and to provide measures quantifying redundant and synergistic information about cognitive variables present in pairwise and higher-order functional interactions. Indeed, classical connectivity measures, such as coherence or amplitude-envelope correlation quantify whether signals are coupled, but not whether that coupling carries information about an external cognitive variable — the gap the present framework is designed to close. To do so, we combined Partial Information Decomposition (Williams and Beer, 2010; Wibral et al., 2017; Lizier et al., 2007; Voges et al., 2024) and information dynamics (Lizier et al., 2013; Voges et al., 2024) frameworks with spectral analysis methods for the extraction of functional interaction measures based on amplitude co-modulations and phases relations between pairs of signals. We first validated our approach through simulations based on the Kuramoto and Stuart–Landau models of coupled oscillators. The former is a classical model for studying oscillatory networks, whereas the latter extends it by incorporating both phase and amplitude, providing a minimal system capable of exhibiting subcritical Hopf bifurcations as well as transient and steady-state oscillations (Ponce-Alvarez and Deco, 2024). We then tested the methods on simulations of multiple oscillators coupled according to macaque monkey structural connectivity (Markov et al., 2014), capturing inter-areal interactions through synchronization and amplitude co-modulations. We finally provided proof-of-concept on local field potentials (LFPs) recorded from macaque cortical regions during a working memory task (Salazar et al., 2012). Our study demonstrates that the approach, named Neural Oscillatory Partial Information Decomposition (NeOPID), provides a relevant tool to disentangle information about cognitive variables in neural oscillations into amplitude co-modulation and phase contributions and to quantify redundant and synergistic information in brain relations, from pairwise to higher-order interdependencies.

## MATERIALS AND METHODS

### Neural Oscillatory Partial Information Decomposition (NeOPID)

We propose an approach based on Partial Information Decomposition (PID) framework and spectral analysis to quantify task-related information contained in oscillatory interactions in the frequency domain. The approach can be used to decompose information about task variables contained in amplitude co-modulations or phase differences between oscillatory signals. In addition, the approach can quantify redundant and synergistic information contained in higher-order oscillatory interactions. The approach is specifically designed for the analysis of neural oscillations and the decomposition of task-related information contained in brain interactions. We refer to the approach as Neural Oscillatory Partial Information Decomposition (NeOPID).

#### Partial Information Decomposition for pairwise and higher-order functional neural interactions

The first building block of NeOPID is based on the Partial Information Decomposition (PID) (Williams and Beer, 2010; Wibral et al., 2017) and information dynamics (Lizier et al., 2013; Voges et al., 2024) frameworks. PID provides a mathematical framework for the decomposition of multivariate mutual information between a system of predictors and a target variable into unique, redundant and synergistic information terms (Williams and Beer, 2010; Wibral et al., 2017). As a reminder, a standard measure of the statistical dependence between two random variables is Shannon’s mutual information. As a reminder, mutual information is defined as:

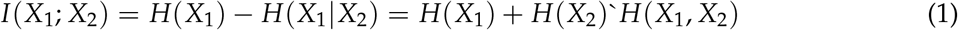

Variables *X*_1_ and *X*_2_ may represent the neural activity of two brain areas or neural populations, respectively. *H*(*X*_1_) is the entropy of *X*_1_, and *H*(*X*_1_|*X*_2_) is the conditional entropy of *X*_1_ given *X*_2_. PID can be used to decompose the information carried by a set of neural signals *X*_1_ and *X*_2_ about a task-related variable *Y, I*(*X*_1_, *X*_2_|*Y*), which may represent the trial-by-trial evolution of stimulus properties, decision or internal variables. When only one signal, *X*_1_ or *X*_2_, individually carries information about a target variable *Y*, this is termed *unique* information - the information that is specific to each source about *Y*. If both *X*_1_ and *X*_2_ convey the same information about *Y*, this shared component is called *redundant* information. Conversely, if neither variable alone provides information about *Y*, but their joint observation does, this corresponds to *synergistic* information. Redundant and synergistic components thus quantify how interactions between neural signals contribute to the information about *Y*. Analytically, the PID for the three-variables case - i.e., two source variables case - decomposes the total mutual information between *X*_1_ and *X*_2_ with a target variable *Y* as:

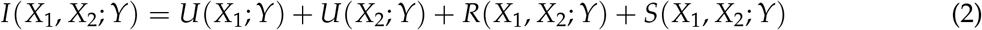

where *U*(*X*_1_; *Y*) and *U*(*X*_2_; *Y*) are the unique information carried by the two areas, respectively, and *R*(*X*_1_, *X*_2_; *Y*) and *S*(*X*_1_, *X*_2_; *Y*) are the redundancy and synergy terms, respectively. In addition, the PID formulation links to classical Shannon measures of mutual information as:

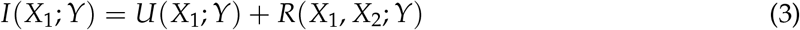

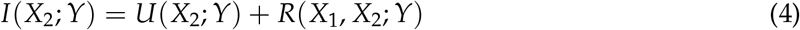

The governing equations form an under-determined system: only three quantities can be computed from the data (i.e., the mutual information quantities *I*(*X*_1_, *X*_2_; *Y*), *I*(*X*_1_; *Y*), and *I*(*X*_2_; *Y*)) for the four terms of the PID (i.e., two unique information terms, redundancy, and synergy). Here, we exploit the so-called *minimum mutual information* (MMI) PID, which has been shown to provide correct estimations for a broad class of systems following a multivariate Gaussian distribution (Barrett, 2015). According to the MMI PID, redundant information carried by pairs of brain regions is given by the minimum of the information provided by each individual source to the target:

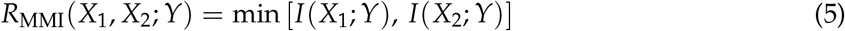

Then, synergistic information can be computed by substituting Eqs. 3, 4, and 5 into Eq. 2 and rearranging the terms:

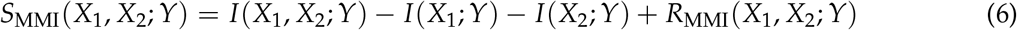

By definition of the MMI approach, redundant functional interactions contain the same information about the target variables as that contained in individual brain areas. On the other hand, synergistic connectivity reveals functional interactions that cannot be explained by individual contributions, but only by their combined and collective coordination.

The approach can be generalized to quantify task-related information present in interactions beyond pairwise relations, the so-called higher-order functional interactions (HOI) or correlations. The definitions of redundancy and synergy can be generalized to higher order by simply iterating over all regions of a multiplet. By considering a multiplet as a set of *n* variables *X*_*n*_ = *{X*_1_, … , *X*_*n*_*}*, the redundant information contained in each higher-order multiplet about the target

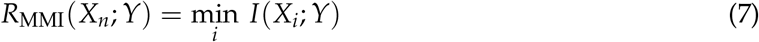

Then, higher-order synergistic information can be computed as:

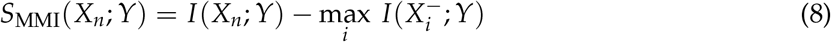

where 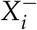 is the set of variables of *X*_*n*_ excluding the source variable *i*. Equations 7 and 8 quantify the amount of redundant and synergistic information about the target variable *Y* contained in higher-order functional interactions between variables *X*_*n*_, respectively. In previous studies, variables *X*_*n*_ have been associated with the time series of nodes in a network (e.g., brain areas) (Combrisson et al., 2025). In such cases, the term higher-order functional interaction reflects the multivariate coupling between brain areas, beyond pairwise relations. Synergistic information (Eq. 8) reflects the extra amount of information about the target variable *Y* carried by the higher-order interactions between brain areas, beyond what is encoded by lower orders. For example, the presence of significant synergistic information about a target variable carried by three brain areas (i.e., a triplet) quantifies the additional information beyond that contained in all pairwise relationships. It therefore reflects a quantity that is fundamentally different from a pairwise metric. In contrast, redundant information does not represent a fundamentally different quantity with respect to lower-order interactions, as it is defined as the minimum of the lower-order marginal mutual information (Eq. 7). The concept of variable *X*_*n*_ is, nevertheless, general, and it can be applied to different metrics. In the current study, the variables *X*_*n*_ are edge-level quantities, such as amplitude co-modulations (Eq. 9) and/or phase-difference variables (Eq. 10), which are each associated with a pair of signals, rather than a single time series. Accordingly, we define interactions between two pairs of edges as higher-order, because they reflect the interaction between more than two nodes. Synergy therefore contains information beyond pairwise relations. These notions and metrics will be clearly described in the section entitled “Partial Information Decomposition of spectral measures for pairwise and higher-order interactions”.

We used Gaussian-Copula Mutual Information (GCMI) (Ince, 2017), which is a semi-parametric binning-free technique to calculate mutual information and entropy estimates. In the GCMI approach, which includes a rank based normalisation of the data, entropy values are computed using the differential entropy formula for a Gaussian distribution. The transformed data allows entropy estimation based on the determinant of the covariance matrix and the assumption of Gaussian marginals. The GCMI is a robust rank-based approach that allows detecting any type of monotonic relation between variables. More precisely, the GCMI is a lower-bound approximate estimator of mutual information for continuous signals based on a Gaussian copula, and it is of practical importance for the analysis of short and potentially multivariate neural signals. Note, however, that the GCMI does not detect non-monotonic (e.g., parabolic) relations. As an example, this methodology has been applied to investigate the information about learning signals, such as information gain, contained in cortico-cortical co-modulations in broadband high-gamma MEG activity (60-120Hz) (Combrisson et al., 2025).

#### Frequency-domain measures of amplitude and phase relations between neural oscillations

The second building block of NeOPID is represented by spectral analyses tools that measure amplitude and phase relations between neural oscillations. Although consensus has not been reached regarding the mechanisms support neural interactions, most accounts emphasize a central role of co-modulations in aperiodic transient dynamics (Vinck et al., 2023, 2025) and phase relations between neural oscillations (Engel et al., 2001; Fries, 2015; Vezoli et al., 2021; Bastos et al., 2015). Decomposing the information contained in phase relations and amplitude co-modulations is therefore crucial.

We consider the cross-spectral density between two zero-mean time series *x*_*n*_(*t*) and *x*_*m*_(*t*), with Fourier transforms *X*_*n*_(*f*) and *X*_*m*_(*f*). The cross-spectrum between two time-domain signals with Fourier transforms *X*_*n*_(*f*) and *X*_*m*_(*f*) is given by 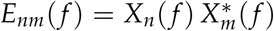, where ∗ represents the complex conjugate. The cross-spectrum is a complex-valued function that can be expressed in polar form as 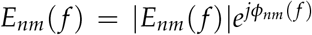. The magnitude |*E*_*nm*_(*f*)| quantifies the strength of frequency-specific linear dependence between the two signals, reflecting the degree of covariance in their oscillatory components at frequency *f* .

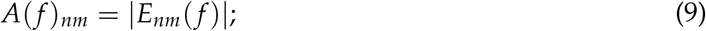

We refer to *A*_*nm*_(*f*) throughout the manuscript as the *amplitude co-modulation* between signals *x*_*n*_ and *x*_*m*_, to distinguish it explicitly from the amplitude (or power) of a single oscillatory signal considered in isolation: it is a pairwise, cross-spectral quantity that co-varies with the joint envelope dynamics of the two signals, not a property of either signal alone. We adopt *amplitude co-modulation* as the default term for this quantity throughout the manuscript.

The instantaneous phase difference at frequency *f* between two oscillatory signals was computed as a symmetry-invariant angular distance derived from the cross-spectra as,

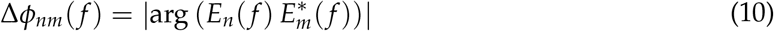

This quantity takes values in [0, *π*]. The use of the absolute value enforces invariance to sign, such that phase offsets of +*θ* and −*θ* are mapped to the same value. This corresponds to a modeling choice in which coupling strength is treated as directionally symmetric, while phase lead–lag information is not retained. Under this representation, a value of 0 corresponds to perfect in-phase alignment, values near *π*/2 correspond to quadrature relationships, and values near *π* correspond to anti-phase alignment. This provides a phase-based distance metric between oscillatory processes that can be used as input for subsequent information-theoretic analyses. Importantly, this measure does not correspond to spectral coherence in the strict statistical sense, but rather to an instantaneous phase distance derived from complex analytic signals. The use of absolute phase differences avoids cancellation effects that arise when signed phase differences are averaged over time in non-stationary or multimodal phase distributions, and is consistent with treating phase as a circular variable in circular statistics frameworks. In other words, we used the absolute phase difference (wrapped to [0, π]), corresponding to the absolute angular distance (Vinck et al., 2010). This avoids cancellation effects inherent to signed phase differences when averaged over time, a known issue in phase-synchronization analysis (Lachaux et al., 1999; Vinck et al., 2010), and is consistent with treating phase as a circular variable (Fisher, 1995). We should also note that such implemented can establish whether task information is present in a symmetric phase relationship, but cannot on its own determine which area leads or drives the other.

#### Partial Information Decomposition of spectral measures for pairwise and higher-order interactions

We then combined spectral measures with PID to provide a decomposition of the total mutual information contained in amplitude co-modulations *A*_*nm*_ and phase difference Δ*θ*_*nm*_ between pairs of brain signals about a task variable *Y* as *I*(*A*_*nm*_, Δ*θ*_*nm*_; *Y*). The mutual information *I*(*A*_*nm*_, Δ*θ*_*nm*_; *Y*), quantifies how much information about the task variable *Y* is jointly carried by the amplitude co-modulations *A*_*nm*_ and the phase differences Δ*θ*_*nm*_ between pairs of signals. In other words, it measures the extent to which task-related changes can be explained by variations in both the shared amplitude co-modulation and the relative phase relationships of the interacting signals across frequencies. The quantity *I*(*A*_*nm*_, Δ*θ*_*nm*_; *Y*) can be interpreted as a sort of “task-related coherence” measure, as it quantifies the mutual information between task variable *Y* and the joint amplitude co-modulations and phase relationships between signals *x*_*n*_ and *x*_*m*_. We use the term “task-related coherence” here in an analogical sense only: despite the name, this quantity is a mutual-information measure between task variable *Y* and the joint amplitude/phase relationship between two signals, and it should not be equated with spectral coherence in the classical, cross-spectral sense. It thus reflects how task variable modulates the spectral coupling between pairs of signals. Task-related coherence, however, conflates contributions from *A*_*nm*_ and Δ*θ*_*nm*_. Here, we propose a decomposition based on PID as described above, yielding unique information contained in either amplitude co-modulations *A*_*nm*_ and Δ*θ*_*nm*_, their redundant and synergistic contributions. The redundant information reflects information about the task variable shared by both amplitude co-modulations and phase differences. Whereas, the synergistic atom represents information about the task variable that can be learned by looking at the interaction between *A*_*nm*_ and Δ*θ*_*nm*_. These terms can be computed using the formalism described above.

These measures can be generalized to quantify the information about task variables contained higher-order interactions by studying multiple interacting edges (i.e., pairs of nodes) via phase differences. The rationale is that pairs of brain regions that share information about task variables through modulations in phase difference may form a network of interacting edges. Such network may track collectively either by means of redundant or synergistic information. This may be measures by considering equations 11 and 12, and using as source variables of the PID the edge-wise phase differences. By considering a multiplet as a set of *k* phase differences between pairs of brain signals, 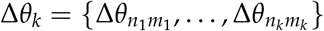, the redundant information contained in each higher-order multiplet about the target variable is given by:

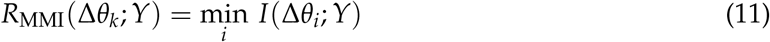

Then, higher-order synergistic information can be computed as:

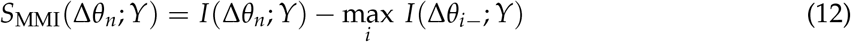

The redundancy and synergy measures quantify how multiple phase-difference interactions between brain signal collectively track information about a task variable. The redundant information *R*_MMI_(Δ*θ*_*k*_; *Y*) captures the portion of task-related information that is shared across all phase-difference pairs within a multiplet, that is, information redundantly represented by each edge. In contrast, the synergistic information *S*_MMI_(Δ*θ*_*n*_; *Y*) reflects information that emerges only when considering the joint activity of multiple edges, beyond what any single edge contributes individually. Together, these measures indicate whether task-related information in a network of interacting edges is organized redundantly (overlapping) or synergistically (complementary) across phase-based interactions.

### Computational model and simulation of neural oscillatory interactions

We tested the NeOPID approach on simulated data using dynamical system models that captures both amplitude and phase-based interactions. We detail the computational model used to simulate the oscillatory dynamics of the nodes, namely the Kuramoto and the Stuart-Landau model. Next, we describe two network configurations used: the first consisting of two interconnected nodes, and the second based on the macaque anatomical connectivity obtained via retrograde tracing, providing the structural connectivity among nodes.

#### The Kuramoto coupled oscillator model

The Kuramoto model describes a set of *N* interconnected nodes with oscillatory dynamics governed by Eq. 13.

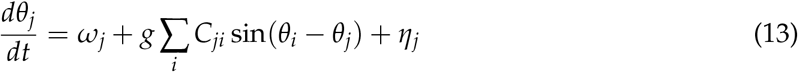

where *ω* is the natural frequency of the oscillator in rad/s, *g* regulates the global coupling between nodes, *C* is the adjacency matrix specifying structural connectivity, and *η* is a white noise input to the system with zero mean and variance *σ*^2^. In our simulations, the temporal resolution of the simulation was set to obtain a central frequency of oscillation of 40Hz. The overall synchrony among nodes is given by the Kuramoto order parameters *r* and *ψ* given by Eq. 14.

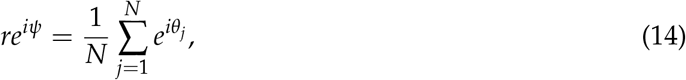

where *r* captures the phase-coherence of the population of oscillators and *ψ* their average phase.

#### The Stuart-Landau network model

Whereas the Kuramoto model describes the phase dynamics of coupled oscillators assuming fixed amplitudes, focusing only on how phases synchronize, the Stuart–Landau (SL) model describes both amplitude and phase dynamics, allowing oscillations to emerge, grow, or decay. Thus, the Kuramoto model is phase-only, while the SL model is a full amplitude–phase oscillator model. The Stuart–Landau model is the canonical normal form of a Hopf bifurcation, which is a fundamental mechanism by which a system transitions from a stable fixed point to oscillatory behavior. In a subcritical Hopf bifurcation, the emerging limit cycle is initially unstable, meaning that small perturbations can cause sudden transitions to large-amplitude oscillations or bistable behavior. In contrast, a supercritical Hopf bifurcation produces a stable limit cycle that grows smoothly from the fixed point as the system parameter changes. The SL model of a single node is given by the Equation 15.

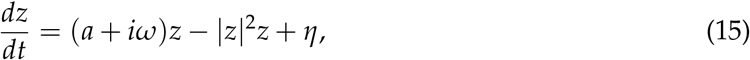

where *z* is a complex-valued variable representing the evolution of the dynamical system, *ω* is the natural frequency of the oscillator in rad/s, *a* is a control parameter dictating how far the system is from the sub-Hopf bifurcation (Figure 1A), |*z*|^2^ is the absolute value of *z*, and *η* is a white noise input to the system with mean zero and variance *σ*^2^. For values of *a* < 0 the system displays damped oscillations; hence, the system *z* = 0 is a spiral or focus point in its state-space. For *a* > 0, the system oscillates, and the stable points are *z* = 0, which correspond to the resting state and 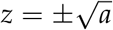 to a limit cycle. The transition between these states happens at the bifurcation at *a* = 0. It is possible to show that the state variable *z* encapsulates the evolution of the amplitude and phase terms of an oscillator. By writing *z* as a vector in the complex plane, with length *r*, and phase *θ*, i.e., *z* = *re*^*iθ*^ and plugging it into Equation 15 with *η* = 0, one gets:

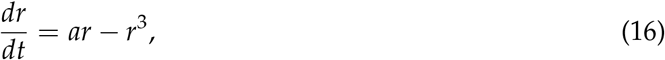

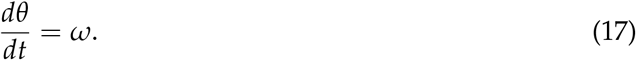

**Figure 1.**
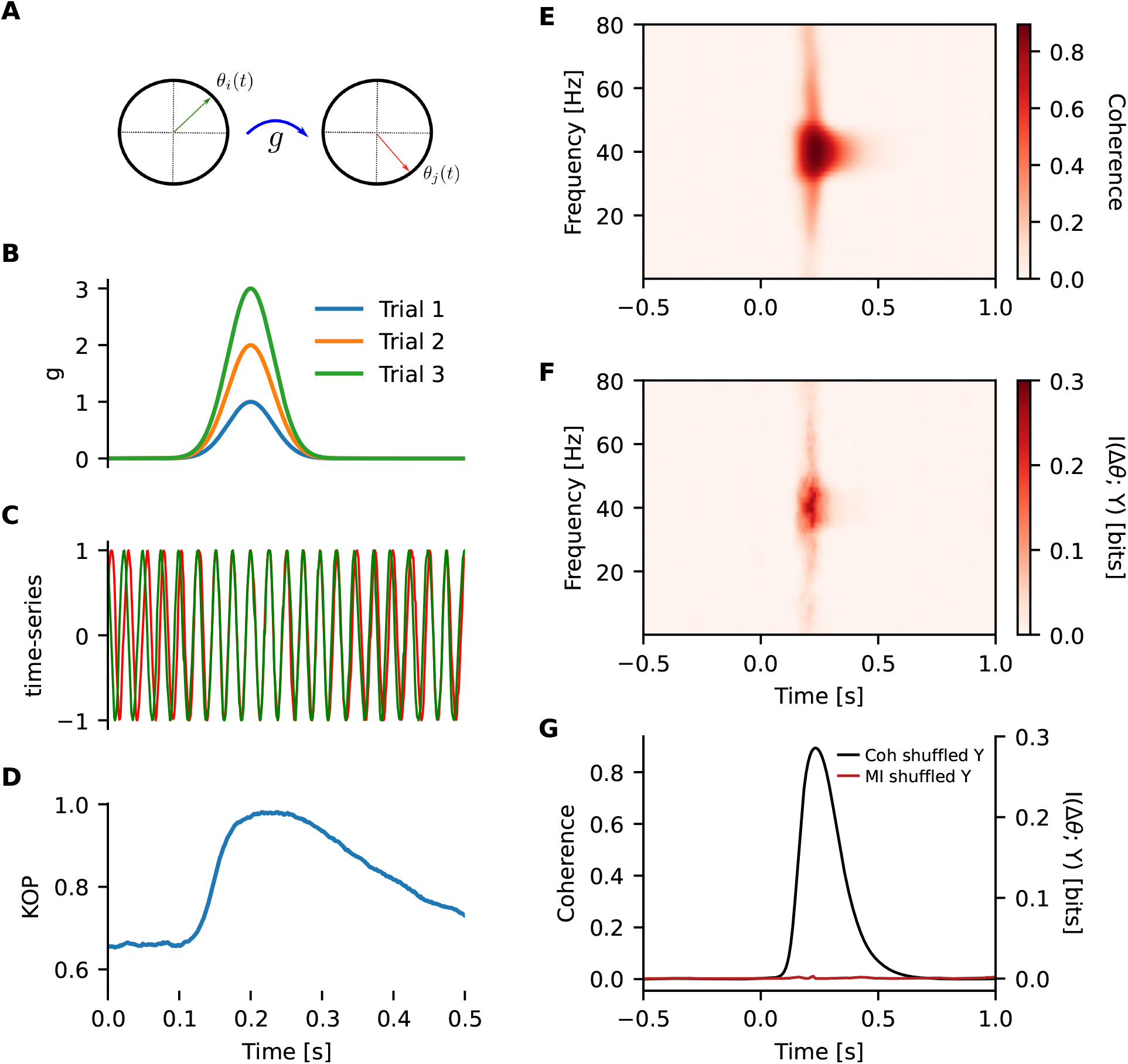
Phase-based encoding in pairwise neural interactions. (**A**) Simulation of two Kuramoto coupled oscillators (see Eq. 13; using identical parameters for both nodes: *ω* = 2*π*40 Hz, *η* = 2.5, and coupling strength *C*_12_ = 10) coupled via an undirected edge. The time series of the simulated dynamical system are represented by their instantaneous phases *θ*(*t*) (depicted by green and red arrows in the unit circle). The coupling strength *g*(*t*) is unidirectional from *X*_*i*_ to *X*_*j*_. (**B**) The coupling strength *g*(*t*) between the two nodes was time-modulated such that it was transiently non-zero from 0.1 to 0.3 s, following *g*(*t*) = 8*Y*/(2*π f*), where *Y* was the maximum coupling amplitude, increased from 1 to 100 across trials (*n* = 1000). (**C**) Time series of the simulated Kuramoto nodes for the trial with the strongest coupling. (**D**) Kuramoto order parameter (KOP) time series averaged across trials. (**E**) Time-frequency coherence spectrogram between the two nodes computed using wavelet decomposition (*n*_cycles_ = *f*_*c*_/4; for *f*_*c*_ ∈ [0.1, 80] Hz). (**F**) Mutual information between the phase difference of the two nodes (computed from the wavelet coefficients as described in the methods section) and the coupling amplitudes *Y*. (**G**) Trial-shuffle control at the frequency of peak coupling (40 Hz). Coherence (black curve) pools cross- and auto-spectra across trials without referencing *Y*, and is therefore identical whether *Y* is left in its real trial order or randomly shuffled. Mutual information (red curve) collapses toward its finite-sample floor once the true trial-to-trial correspondence between Δ*θ* and *Y* is broken, indicating that coherence is insensitive to a task variable that NeOPID captures.

Notice in the differential equations 9 and 10, amplitude and phase components evolve independently. The fact that the amplitude *r* and phase *θ* terms evolve in parallel makes it convenient for studying how cognitive variables are reflected in amplitude co-modulations or phase relations.

#### Whole brain cortical model based on Stuart-Landau oscillators

The Stuart Landau model can be used to simulated whole-brain oscillatory networks by coupling nodes via an adjacency matrix *C*_*ij*_ representing the structural connectivity between nodes.

To determine the inter-regional connectome and the number of projections originating from each cortical area in the whole-brain SL model, we used data reported by Markov et al. (2014), encompassing 29 areas from the macaque cerebral cortex. Restricted to these 29 areas, the connectome is edge-complete, i.e. there are no missing inter-regional connections that have not been characterized (Markov et al., 2013). The directed weight from a source to a target area is estimated to be proportional to the Fraction of Labeled Neurons (FLN). The FLN is defined as the number of labeled neurons in a source area divided by the total number of labeled neurons in all source areas. As retrograde tracers are used, the FLN can be considered as an indication of the relative of neurons sending long-range projections from one source area to a target.

The Equation 15 of the full network model is thus given by:

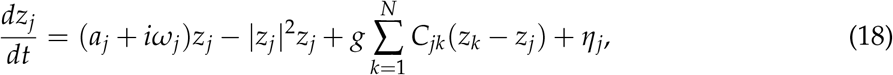

where the indexes *j* and *k* (notice that each node is theoretically allowed a distinct intrinsic frequency and *a* parameter Ponce-Alvarez and Deco (2024)), whereas *g* is a parameter regulating the global coupling between each node in the network. The coupling strengths are scaled with the hierarchy (*h*) of each cortical area to amplify inputs arriving to associative areas, yielding:

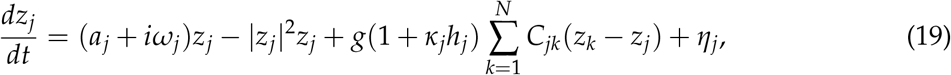

where, *h*_*j*_ is the hierarchical level of node *j*, and *κ*_*j*_ a parameter regulating the amplification due to hierarchy.

#### Simulating task-related modulations

Simulation of task-related modulation of the network dynamics and interactions to the Kuramoto and Stuart-Landau simulations was done by modulating the global coupling among nodes. That is, the parameter *g* in Eqs. 13 and 15 was treated as a dynamic variable *g* = *g*(*t*). More specifically, the coupling strength among nodes was transiently modulated from 0 to 0.4 ms according to *g*(*t*) = 8*Y*/(2*π f*), where *Y* was the maximum coupling amplitude which was increased from 1 to 100 across trials. When computing information-theory quantities the maximum coupling amplitudes *Y* was considered as the continuous stimulus label.

### Eletrophysiological dataset: distributed neural oscillations during working memory

In order to test our approach on neural data, we analyzed local field potentials (LFPs) recorded simultaneously from the lateral prefrontal and posterior parietal cortical areas, exhibiting content-specific coherence in the beta band (15–30 Hz) during working memory tasks in macaque monkeys (Salazar et al., 2012). Monkeys were trained to perform an oculomotor delayed match-to-sample task, which required them to match either the identity or the position of the sample object. During visual fixation, a sample stimulus, consisting of one of three possible objects, was presented for 0.5 s, followed by a random delay of 0.8–1.2 s. At the end of the delay, a match stimulus was presented, consisting of the previous sample object (target) and a distractor object at a different position. A saccadic eye movement to the target was rewarded with juice.

As a proof of concept, we analyzed a single representative recording session, comprising LFPs recorded simultaneously from several prefrontal and posterior parietal sites. Trials were retained for analysis only if they jointly met three quality criteria: the monkey produced a valid behavioral response, the recording was free of major artifacts, and the trial passed the quality control applied at acquisition; trials failing any of these criteria were excluded. The identity-matching and location-matching versions of the task were interleaved within the session, so the task-relevant stimulus feature varied from trial to trial: on identity-matching trials, the label used for the analysis was the identity of the sample object, whereas on location-matching trials, it was the sample object’s spatial position. This trial-wise selection defines the discrete task (or cognitive) variable *Y* used throughout the pairwise and higher-order analyses of this dataset, so that on every trial *Y* corresponds to the specific stimulus feature the animal was required to report.

For each retained trial, LFPs were epoched from −0.4 to 1.7 s relative to sample onset, spanning the pre-stimulus baseline, sample presentation, and part of the delay period. Time-frequency decomposition was performed with a multitaper approach (Slepian tapers; time-bandwidth product of 5, corresponding to approximately nine tapers) over 31 frequencies linearly spaced between 5 and 35 Hz, covering the theta, alpha, and beta bands. The number of cycles per wavelet scaled with frequency to keep a comparable time-frequency resolution trade-off across the band. For every pair of recording sites, the cross-spectrum was obtained by averaging the product of one site’s complex time-frequency coefficients with the complex conjugate of the other’s across tapers, and decomposed into an amplitude co-modulation term and an (unsigned) phase-difference term, following the same definitions used throughout the manuscript (Eqs. 9 and 10). Mutual information between these features and the trial-wise stimulus label *Y* was estimated with the Gaussian-copula estimator implemented in the Frites package (Combrisson et al., 2022), and the statistical significance of the resulting time-frequency maps was assessed against a null distribution obtained by repeatedly permuting the trial-to-stimulus-label correspondence while leaving the recorded activity itself unchanged (1000 permutations for the pairwise analyses in Fig. 6, 200 for the higher-order analyses in Fig. 7 given their higher computational cost), with cluster-based correction for multiple comparisons across time and frequency.

### Available code resources

All models used for numerical simulations were implemented in Python, relying on scientific computing libraries such as MNE-python, NumPy, SciPy, and JAX. The functions employed for information-theoretic and partial information decomposition (PID) analyses are available in the

Frites package (Combrisson et al., 2022) and the HOI package (Neri et al., 2024). The dependencies required to run the simulations, along with five Jupyter notebooks containing step-by-step code to reproduce the figures presented in this manuscript, are provided in the following GitHub repository https://github.com/brainets/neopid.

## RESULTS

### Overview of the NeOPID approach

We present a novel information decomposition approach for brain oscillatory networks, named Neural Oscillatory Partial Information Decomposition (NeOPID). NeOPID quantifies information about cognitive and task variables, such as sensory, decision or cognitive factors, contained in oscillatory interactions by combining partial information decomposition framework (Williams and Beer, 2010; Wibral et al., 2017) with spectral analyses tools and functional interaction measures. NeOPID addresses two open questions. First, how can we quantify task-relevant information contained in phase relations, amplitude co-modulations, or both, and how these two coding channels interact? Second, is information organized redundantly or synergistically across interacting brain regions, from pairwise to higher-order interactions? In other words, NeOPID separates information contained in amplitude co-modulations and phase differences between neural signals in the frequency domain, and it disentangles redundant and synergistic information present in pairwise and higher-order relations. The pipeline is composed of different steps: i) it considers neural oscillations at each recording site; ii) it performs cross-spectral decomposition into frequency-resolved amplitude co-modulation and (unsigned) phase-difference features between pairs, or multiplets, of site; iii) it performs multivariate information decomposition analysis between spectral metrics and the task/cognitive variable via PID, so to extract unique, redundant and synergistic components. Unlike recent information-theoretic approaches which characterize signal-to-signal interactions, such as those introducing frequency-specific PID of oscillatory interactions (Faes et al., 2021; Antonacci et al., 2025) and O-information rate (Faes et al., 2022; Sparacino et al., 2023), NeOPID quantifies information about external task variables encoded in such interactions. We tested the methodology on ground-truth neural interactions generated using oscillatory network models encoding task variables via pairwise and higher-order interactions. We then tested the methods on oscillatory activity generated using dynamical models based on anatomical connectivity from non-human primates (Markov et al., 2014). Finally, we provide a proof-of-concept on simultaneously-recorded local field potentials (LFPs) recorded from the fronto-parietal network of macaque monkeys performing a working memory task (Salazar et al., 2012). We provide a detailed description of the results obtained from simulation and data analysis studies in the following sections.

### Information decomposition about cognitive variables in pairwise phase relations

We simulated the modulation of neuronal interactions through changes in neuronal synchronization and coupling strength in a coupled two-node model (pairwise interaction). This may correspond, for example, to a top-down attentional effect modulating the functional connectivity between two brain regions (e.g., V1 and V4) by transiently improving phase synchrony through task-specific increases in coupling strength (Fries, 2005; Buschman and Miller, 2007). The modulation in phase synchrony would parallel the strength of attentional modulation, enabling phase-based information about cognitive variables (Womelsdorf et al., 2007).

We simulated a two-node Kuramoto model with a central oscillation frequency of 40 Hz and unidirectional coupling from *X*_*i*_ to *X*_*j*_. As a reminder, the Kuramoto model describes the phase dynamics of coupled oscillators assuming fixed amplitudes, it therefore allows only phase encoding. To simulate a task-related modulatory effect of the phase relationship between the two nodes both in time and trials, we varied the coupling strength *g*(*t*) (see Methods). Within trial, the coupling strength followed a Gaussian profile (Fig. 1B), producing transient phase synchronization in the time interval from 0.1 to 0.3 s. Across trials, the amplitude of the Gaussian profile was varied from 1 to 100 (over *n* = 1000 trials), thereby producing an encoding of the task-related variable *Y* into the phase coupling between *X*_*i*_ and *X*_*j*_. Panel 1C shows exemplar time series for the two nodes, illustrating the clear synchronization during the stimulus interval. Consistently, the Kuramoto order parameter (KOP) exhibited a transient increase during the input period (Fig. 1D).

Time-frequency representation of spectral coherence between nodes was computed using wavelet transform methods, revealing a pronounced transient synchronization at 40 Hz (Fig. 1E). We then computed the mutual information in the time and frequency domains between the phase difference of *X*_1_ and *X*_2_ (Δ*θ*) and the task variable *Y*, which reflected the across-trial modulation of coupling amplitude. The resulting spectrum showed a clear peak at 40 Hz, demonstrating that the modulation of coupling strength was contained in the phase relationship between the two nodes (Fig. 1F). Overall, the results provide evidence that the proposed methodology successfully captures the amount of information contained in phase relations between nodes about a task-related variable modulating coupling strength.

In order to show that coherence (Fig. 1E) and *I*(Δ*θ*; *Y*) (Fig. 1F) reflect different properties of the data, we performed a trial-shuffle control at the frequency of peak coupling (40 Hz). Coherence and mutual information were each recomputed after randomly shuffling the trial-to-trial correspondence between the coupling trajectory and *Y*. Since coherence is computed by pooling cross- and auto-spectra across trials without referencing *Y*, it was unaffected by the shuffle (Fig. 1G, black). Mutual information, in contrast, collapsed toward its finite-sample floor once the shuffle disrupted the true trial-to-trial correspondence (Fig. 1G, dashed red), confirming that the modulation observed in Panel F reflects task-related information. These results illustrate the distinction between classical functional connectivity measures, which quantify whether two signals are coupled, and NeOPID, which allow the distinction between intrinsic (ongoing) and extrinsic (task-related) coupling between neural oscillations, either via phase relations and/or amplitude co-modulations.

### Information about cognitive variables in higher-order interactions via phase relations

We then evaluated NeOPID’s ability in quantifying the information contained in higher-order oscillatory interactions via phase-only relations. The rationale was to generalize the previous analysis to account for mutual information between two or more phase differences and the task variable *Y*. To do so, we employed the Partial Information Decomposition (PID) framework, which provides a mathematical formalism for the decomposition of multivariate mutual information between a set of predictors and a target variable into unique, redundant and synergistic information terms (Williams and Beer, 2010; Wibral et al., 2017). In the current study, the PID formalism was used to decompose the total mutual information between two phase differences between nodes Δ*θ*_1_ and Δ*θ*_2_, respectively, and the task variable modulating coupling strength *Y*, into redundancy and synergy terms. The redundant information captures the portion of task-related information that is shared across all phase-difference pairs, that is, information redundantly represented across edges. In order words, redundant information reflects the minimum amount of coupling that is present across all edges. It therefore reflects a higher-order measure that correspond to the minimum pairwise coupling with the task variable. In contrast, synergistic information reflects information that emerges only when considering the joint activity of two edges, beyond what any single edge contributes individually. Synergistic information is, by construction, absent from any marginal or pairwise metric. Recovering synergy requires a genuinely multivariate readout beyond pairwise information. Together, these measures indicate whether task-related information in a network of interacting edges can be quantified via redundant (shared) or synergistic (complementary) information in higher-order phase relations.

We simulated modulation of neuronal interactions through changes in neuronal synchronization and coupling strength in a network with three nodes using Kuramoto coupled oscillators. Generally, a chain structure promotes redundant encoding, because information propagates sequentially, causing downstream nodes to inherit overlapping representations from upstream sources, whereas synergistic encoding requires convergent integration (i.e., colliders) of independent inputs. To simulate a higher-order oscillatory interaction in which phase relations contain task-related information with a greater redundant (shared) than synergistic component, we therefore simulated a feedforward Kuramoto model organized in a chain structure (Fig. 2A). The coupling strength varied across trials, and in a trial-based and time-varying manner across all edges. As expected, the time-frequency representation for the chain of oscillators showed stronger redundant than synergistic information (Fig. 2B). We then simulated a higher-order oscillatory interaction dominated by synergistic encoding using a collider structure (Fig. 2C). This structure produced a relatively stronger synergistic information as evidenced by the peak value in the time-frequency representation (Fig. 2d). Finally, we simulated the oscillatory activity of a network of three visual cortical areas (V1, V2 and V4 circuit) connected anatomically accordingly to retrograde tracing data, the so-called fraction of labeled neurons (FLN) (Markov et al., 2013). Within such recurrent structure, task information produced comparable redundant and synergistic information contained in higher-order phase relations (Fig. 2F). To conclude, our simulations demonstrate that redundant and synergistic information about cognitive variables can be disentangled for higher-order interactions, therefore opening the possibility to study the distributed encoding of cognitive variables in oscillatory networks via phase relations.

**Figure 2.**
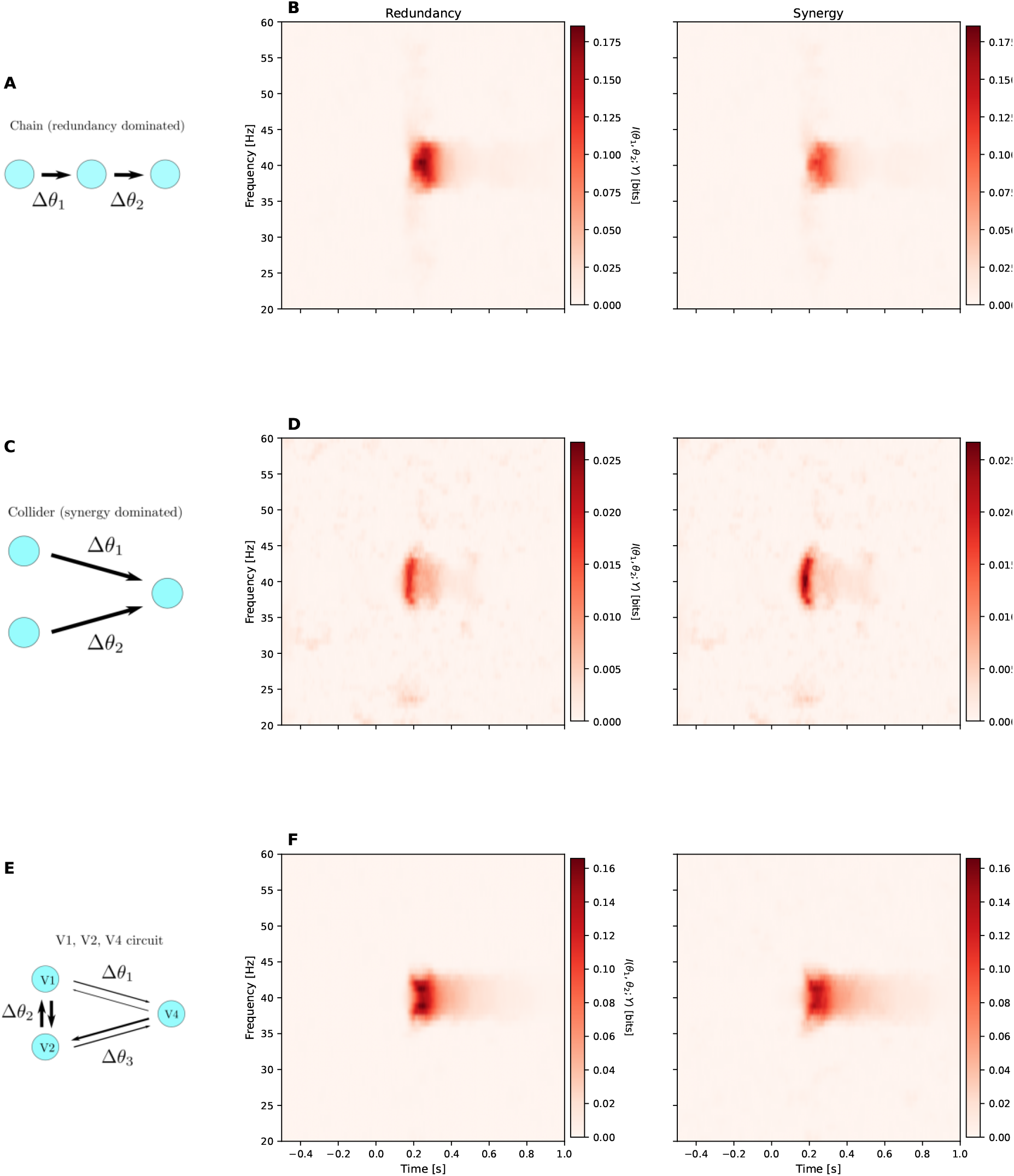
Phase-based information in higher-order interactions among phase-difference (edge) variables. Simulation of three Kuramoto coupled oscillators (Eq. 13) with the same parameters for all nodes (*ω* = 2*π* 40 Hz; *η* = 1.5), coupled via unidirectional edges. The coupling strength among nodes was transiently modulated from 0.1 to 0.3s using a Gaussian profile with coupling amplitude increasing from 1 to 100 across trials (*n* = 1000). (**A**) Redundant encoding was simulated using a chain structure. Phase differences between nodes Δ*θ*_1_ and Δ*θ*_2_ were derived from the Wavelet coefficients of the nodes’ time-series (Eq. 10; *n*_cycles_ = *f*_*c*_/2; for *f*_*c*_ ∈ [20, 60] Hz). (**C**) Synergistic encoding was simulated using a collider structure. (**E**) Simulation of a three-nodes Kuramoto model structured in a recurrent connectivity. Structural connectivity strengths are proportional to their anatomical connectivity measured via retrograde tracing (FLN; Markov et al. (2013)) between three hypothetical visual cortical areas V1, V2 and V4 circuit. The effective connectivity between nodes *i* and *j* is given by *C*_*ij*_ *FLN*_*ij*_, and the arrows representing edges are scaled accordingly. (**B, D and F**) Time–frequency representation of redundant (left panel) and synergistic (right panel) information contained in phase differences Δ*θ*_1_ and Δ*θ*_2_ about the task-related variable *Y*, computed using the PID approach (Eqs. 5 and 6).

### Dissociating information about cognitive variables in phase relations and amplitude co-modulations in oscillatory networks

Recent hypotheses suggest that aperiodic transients and oscillatory dynamics may differentially contribute to neural interactions along feedforward and feedback pathways for sensory inference and predictive processing (Vinck et al., 2023, 2025). Within this framework, neural oscillations are proposed to contain and stabilize neural representations over time and facilitate plasticity processes during the late, feedback-dominated phases of sensory processing. Transients and non-oscillatory dynamics are thought to constitute the primary mechanism for inter-areal communication during sensory inference. Dissociating the contribution of phase relations of oscillatory dynamics and amplitude co-modulations of transients is therefore important for testing such hypotheses. To investigate how NeOPID can be used to address this issue, we simulated a two-node network based on Stuart-Landau oscillators, instead of Kuramoto model. Indeed, whereas the Kuramoto model describes the phase dynamics of coupled oscillators assuming fixed amplitudes, the Stuart–Landau (SL) model describes both amplitude and phase dynamics, allowing oscillations to emerge, grow, or decay. We simulated task-relevant information by modulating the coupling from *X*_*i*_ to *X*_*j*_ as a Gaussian input of varying amplitude across trials, similar to previous simulations based on Kuramoto coupled oscillators. However, contrary to the Kuramoto model, the SL model does not produce encoding exclusively via phase synchronization, because SL oscillators have both amplitude and phase degrees of freedom. Although the NeOPID approach is generalizable to the frequency domain, we computed the mutual information between the amplitude co-modulation and phase differences between the broadband signals, for the specific application to the Stu-art–Landau model. In fact, the Stuart–Landau model already produces complex-valued signals, so a Hilbert-based transformation to obtain analytical signals is not required. The total mutual information between the joint amplitude co-modulation and phase difference and coupling strength was then decomposed into unique, redundant and synergistic terms using PID as described in the Methods.

We simulated a two-node SL coupled oscillators model with unidirectional coupling *g*(*t*) from *X*_*i*_ to *X*_*j*_ (Fig. 3A and B). The total mutual information between the joint amplitude co-modulation and phase difference with task variable displayed a clear peak that paralleled the modulation in coupling strength (green curve in Fig. 3C). Most of the interdependence was carried by the amplitude co-modulation (orange curve in Fig. 3C). Given that modulation of *g*(*t*) drives both amplitude modulation (AM) and phase modulation (PM) in a SL model, we expected the encoding of coupling strength to produce synergistic and redundant effects. Since our approach allows the decomposition of total mutual information in redundant and synergistic terms, we quantified each contribution separately. Indeed, information decomposition showed that a large component is present synergistically in amplitude co-modulation (orange curve in Fig. 3D), with an additional contribution of redundant information (blue curve in Fig. 3D). Given that most of the mutual information among variables is due to the coupling between amplitude co-modulation, we also observed that the unique information in amplitude co-modulation dominated (orange curve in Fig. 3E), whereas the unique information in phase difference was null (blue curve in Fig. 3E). Overall, the simulation showed that the contribution of amplitude co-modulations and phase relations between oscillatory nodes can be disentangled and quantified in terms of redundant and synergistic contributions.

**Figure 3.**
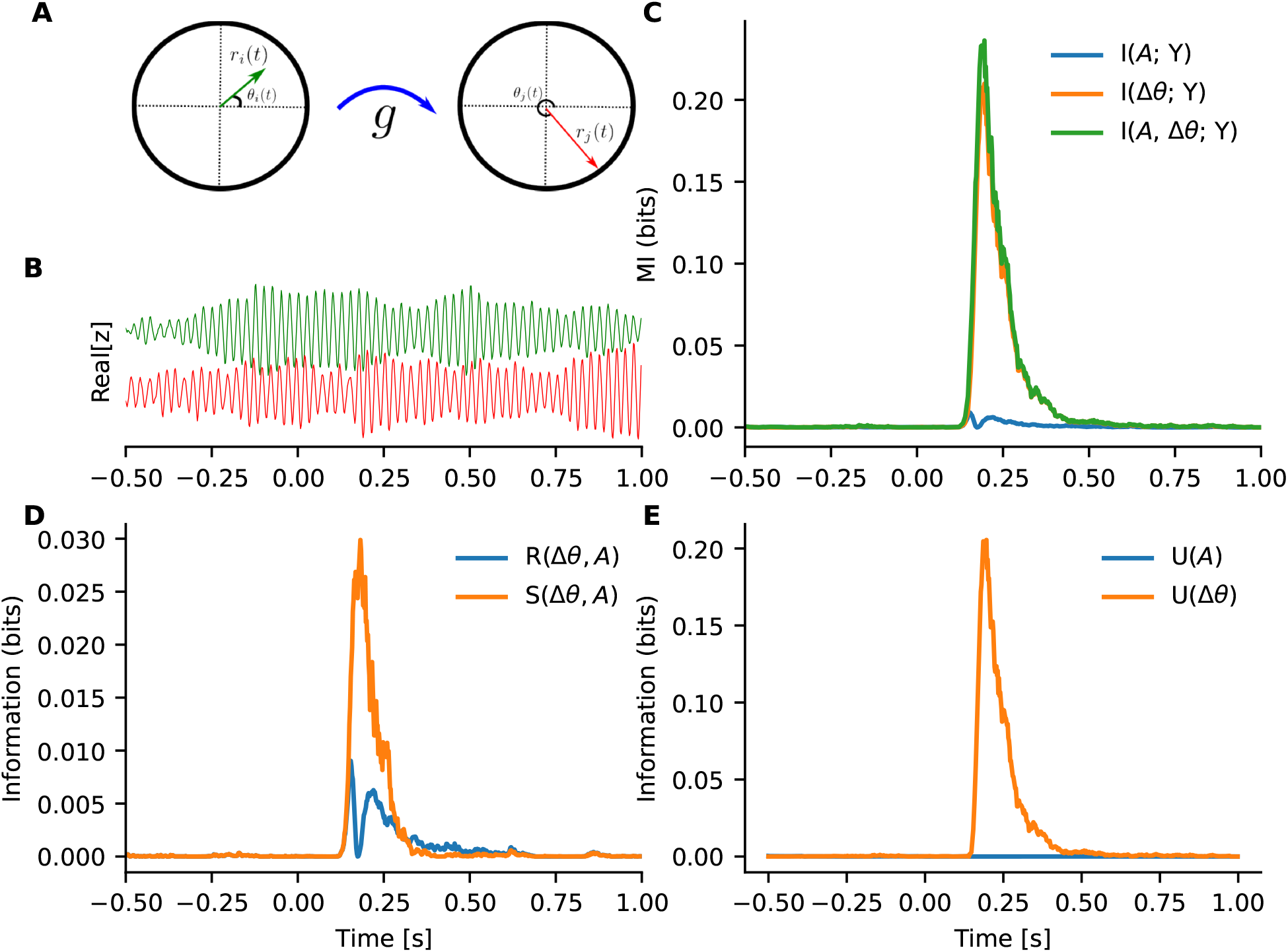
Phase vs. amplitude information in the Hopf model. (**A**) Simulation of two Stuart-Landau oscillators (See Eq. 18; using the same parameters for both nodes *a* =− 5; *ω* = 2*π*40 Hz; *η* = 0.0001; and coupling strength *C*_12_ = 1) coupled via a unidirectional link from *X*_*i*_ to *X*_*j*_. The coupling strength *g* among nodes was transiently modulated from 0 to 0.4 s according to *g*(*t*) = 8*Y*/(2*π f*), where *Y* was the maximum coupling amplitude which was increased from 1 to 100 across trials (*n* = 5000). (**B**) Real part of the (Real(*Z*)) time-series for the simulated SL system. (**C**) Stimulus-related information contained in the broadband joint amplitude co-modulation (blue; 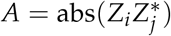); the phase-difference (orange; 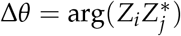), and both (green). (**D**) The redundant (blue) and synergistic (green) information about the stimulus contained in the phase-difference and the joint amplitude co-modulation. (**E**) The unique information about the stimulus in the phase-difference (orange) and the joint amplitude co-modulation (blue).

### Information about cognitive variables in phase relations and amplitude co-modulations in whole-brain cortical network model

We next studied the encoding of task variables in whole-brain cortical network oscillatory models using empirically estimated inter-areal coupling strengths. Our goal was to investigate whether we could quantifying information contained in amplitude co-modulations and phase relations in neural oscillatory networks whose interactions are constrained by anatomical data, and to isolate redundant and synergistic terms for both pairwise and higher-order oscillatory interactions. To do so, we created a model of the macaque cortex by simulating a network of coupled Stuart Landau oscillators. We adopted the approach proposed in Joglekar et al. (2018), assuming that the anatomical connectivity between cortical areas is given by the fraction of labeled neurons (FLN) (Markov et al., 2013), which provides a quantitative map of cortical projections (Fig. 4A). The time series display oscillatory patterns across the nodes of the network and a modulation following modulation of coupling strength in the whole network (Fig. 4B). As a reminder, the modulation of the coupling strength reproduced a task-related dependence of varying amplitude across trials and time, similarly to previous simulations. We first investigated how information about task variable was contained at the level of amplitude co-modulations *A*_*ij*_ and phase differences Δ*θ*_*ij*_ between pairs of nodes. To do so, we focused on two brain areas that are known to be strongly anatomically connected: the primary and secondary visual cortices (areas V1 and V2, respectively). We observed that the total (blue curve) and marginal (green and orange curves) mutual information successfully tracked the dynamics of information encoding (Fig. 4C), thus suggesting that information about cognitive variable was contained in both amplitude co-modulations and phase relations. To quantify the interaction between amplitude co-modulation and phase encoding, we computed redundant and synergistic information. The temporal evolution of redundant and synergistic encoding showed that both contributed to the overall interaction, peaking approximately at the same time (Fig. 4D). We then quantified the unique information in amplitude co-modulations and phases. We observed a positive contribution for both, but peaking at slightly different time points, suggesting that the information in phase synchrony peaks earlier than that in amplitude comodulation (Fig. 4E). As a control analysis, we studied two distant brain regions: the primary visual cortex (area V1) and the anterior cingulate sulcus (area 24c). Although distant, we observed that the total mutual information displayed a strong modulation, therefore suggesting that information about task variable propagates effectively to prefrontal areas (blue curve in Fig. 4F). However, our analysis showed that most of the information was contained in amplitude co-modulations rather than phase relations (orange and green curves in Fig. 4F, respectively). As a confirmation, the redundant and synergistic information in amplitude co-modulations and phases were negligible (Fig. 4G), and most was contained in amplitude co-modulations in a unique way (Fig. 4H).

**Figure 4.**
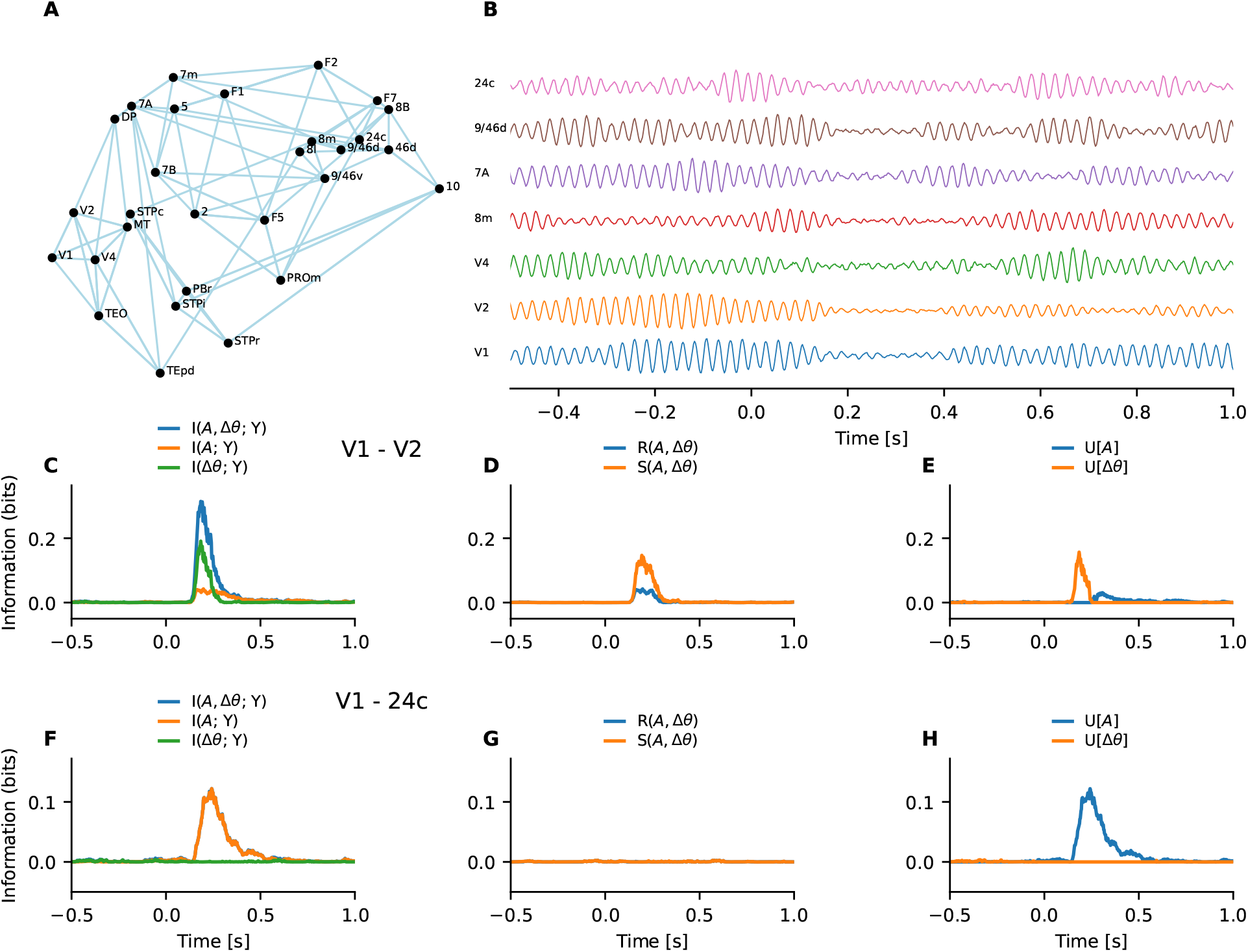
Amplitude co-modulation and phase contributions in the SL whole-brain model. (**A**) Structural connectivity derived from the dataset reported in Markov et al. (2014). Here we show it as a binary network with only the top four connections of each cortical area being drawn (FLN greater than the 87 percentile for each source area). (**B**) The real part of the (Real(*Z*)) time-series for the simulated SL whole-brain model for selected 9 cortical areas in one trial. The black time-series on the bottom show the global coupling dynamics *g*(*t*). The coupling strength *g* among nodes was transiently modulated from 0 to 0.4 s according to *g*(*t*) = 8*Y*/(2*π f*), where *Y* was the maximum coupling amplitude which was increased from 1 to 100 across trials (*n* = 1000). (**C**) The time-series of the total mutual information between V1 and V2 contained in their joint amplitude co-modulation *A* and phase-difference Δ*θ* about the stimulus (blue curve). Only their amplitude co-modulation (orange curve), and only their phase-difference (green). (**D**) The redundancy (blue curve) and synergy (orange curve) between the joint amplitude co-modulation and phase-difference concerning the stimulus. (**E**) The unique stimulus information contained in the joint amplitude co-modulation (orange curve), and only their phase-difference (green). (**F-H**) The same as **C-E** but for areas V1 and 24c. (**I**).

We then generalized the analysis of oscillatory interactions beyond pairwise relations, the so-called higher-order interactions (HOIs). In other words, we investigated whether large-scale oscillatory network encoding task information can recruit more than pairs of nodes. Indeed, we observed oscillatory HOIs among pairs of edges (i.e., pairs of phase-difference variables) displaying redundant and synergistic information about task modulation via phase relations. The results showed that higher-order interactions among two edges, spanning four nodes, contained significant information via coupled phase relations (Fig. 5A and B). This can be interpreted as a correlate of distributed information about cognitive variables contained in collective phase relations. Finally, we investigated higher-order information between three edges. The results showed that several multiplets displayed non-zero redundant and synergistic information (Fig. 5C and D).

**Figure 5.**
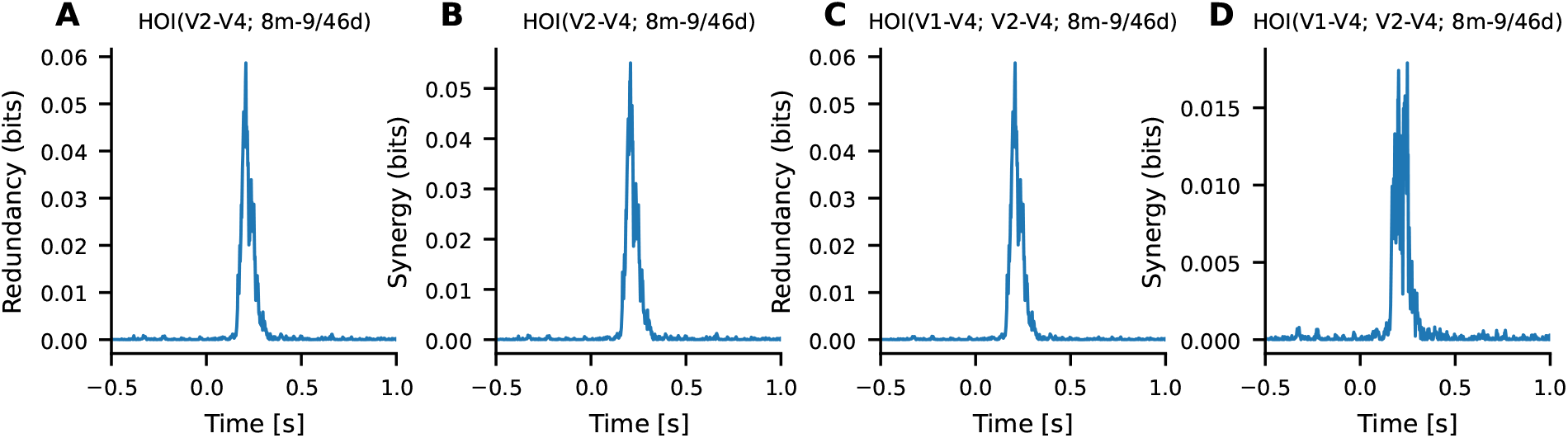
High-order interaction among phase-difference edges in the SL whole-brain model. High-order redundant (A) and synergistic (B) information contained in pairs of edges via phase difference (Δ*θ*_1_ and Δ*θ*_2_, respectively). The PID quantities are computed for the phase-difference time-series between nodes *i*-*j* and *l*-*k*. The curves shown are for the two disconnected edges (four nodes in total) with highest synergy and redundancy formed by the edges V2-V4 with 8m-9/46d (A and B). Oscillatory higher-order redundant (C) and synergistic (D) information contained in three edges (V1-V4, V2-V4 and 8m-9/46d) and five nodes.

Overall, these results show that NeOPID is able to disentangle information about task variables contained in amplitude co-modulations and phase relations between pairs of nodes in a whole-brain oscillatory model. In addition, the results provide evidence that distributed information about cognitive variables via phase relations between nodes beyond pairwise relations can be extracted from oscillatory activity.

#### Distributed information in cortical oscillatory networks during working memory

Finally, we aimed to provide proof-of-concept using electrophysiological data recorded from non-human primates. We analyzed local field potentials (LFPs) simultaneously recorded from lateral prefrontal and posterior parietal cortical areas. Previous studies have shown that LFPs display content-specific beta-band (15–30 Hz) coherence during working memory tasks (Salazar et al., 2012). We first computed the total mutual information between stimulus type and the joint amplitude co-modulation and phase relations for two LFPs located in the caudal part of the inferior parietal lobule (area PG) and area PE (in the dorsal portion of the superior parietal lobule) in the beta band (from 15 to 30 Hz). We observed a stimulus-dependent modulation of total mutual information after stimulus onset and during the delay period of the task (Fig. 6A). The total mutual information was decomposed into unique contributions due to amplitude co-modulation and phase relations. For the session under investigation, the results show that most of the information about stimulus is contained in amplitude co-modulations (Fig. 6B), rather than phase relations (Fig. 6C).

**Figure 6.**
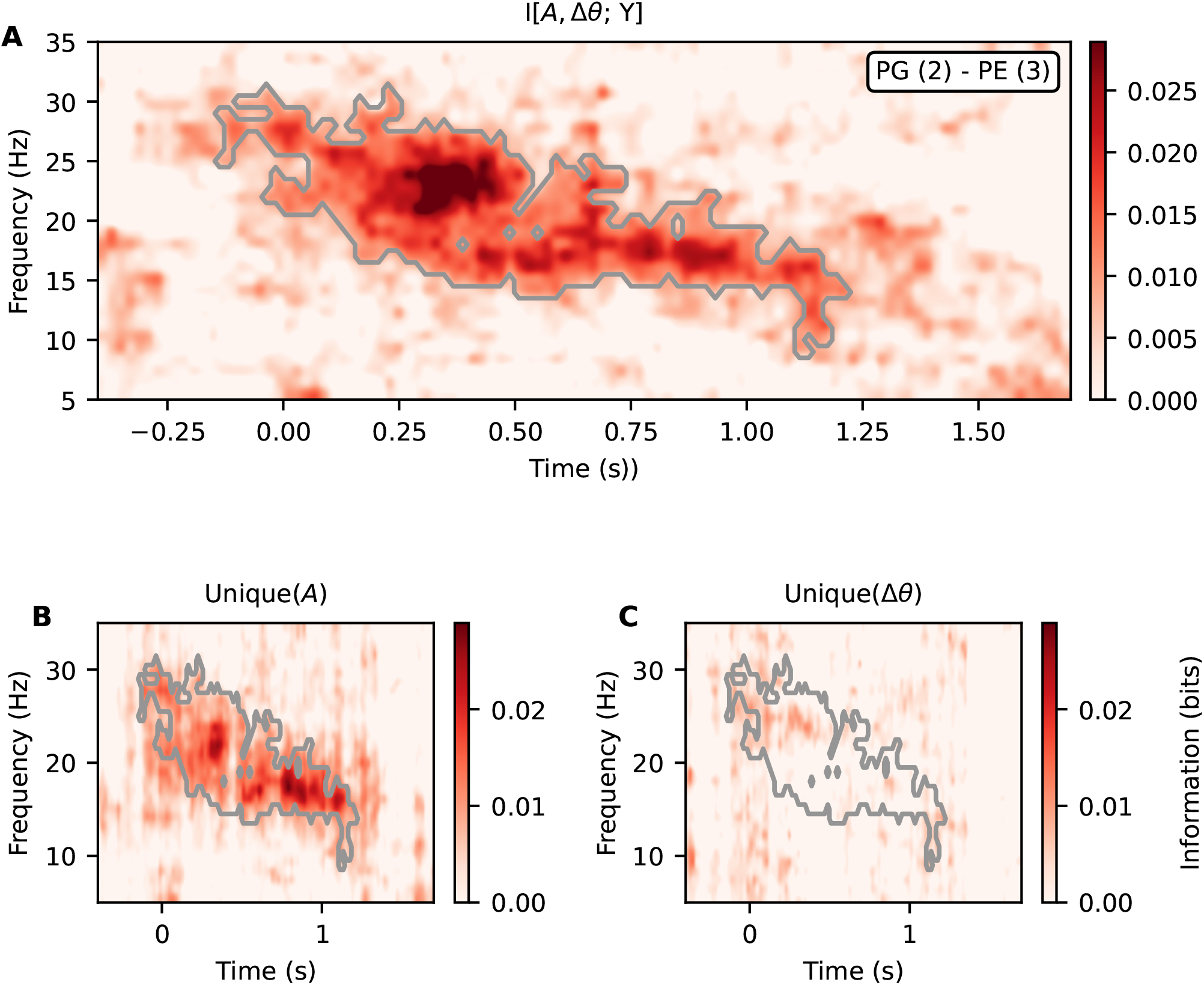
Phase relations and amplitude co-modulations in fronto-parietal recordings during working memory. **A** Stimulus-related coherence given by the total mutual-information between phase relations and amplitude co-modulations about the stimulus (Eq. 2) for two parietal channel pair (PG - PE). **B**,**C** Unique information about the stimulus contained in the joint amplitude co-modulation and phase-difference components of the signal. Areas delimited by the white line correspond to significant effects obtained via shuffling surrogates (*n* = 1000; *p* < 0.05; corrected using cluster statistics).

Next, we investigated the presence of higher-order oscillatory interactions among pairs of edges derived from LFPs. To do so, we analyzed two pairs of LFPs recorded from the fronto-parietal network. The first pair included channels located the PG and PE areas. The second pair included channels located in the dorsomedial frontal cortex area 8B (anterior to the frontal eye fields, area 8) and parietal area PE. (Fig. 7A) shows the total mutual information between amplitude co-modulations and phase relations, and stimulus type, showing clear higher-order information structure. We then investigated whether the observed mutual information was due to redundant or synergistic interactions. The results suggest that both redundant (Fig. 7B) and synergistic (Fig. 7C) interactions between higher-order phase relations are present. Overall, we provided proof-of-concept that NeOPID can be used to disentangle information contained in amplitude co-modulations and phase relations in oscillatory networks, and to identify distributed oscillatory higher-order interactions.

**Figure 7.**
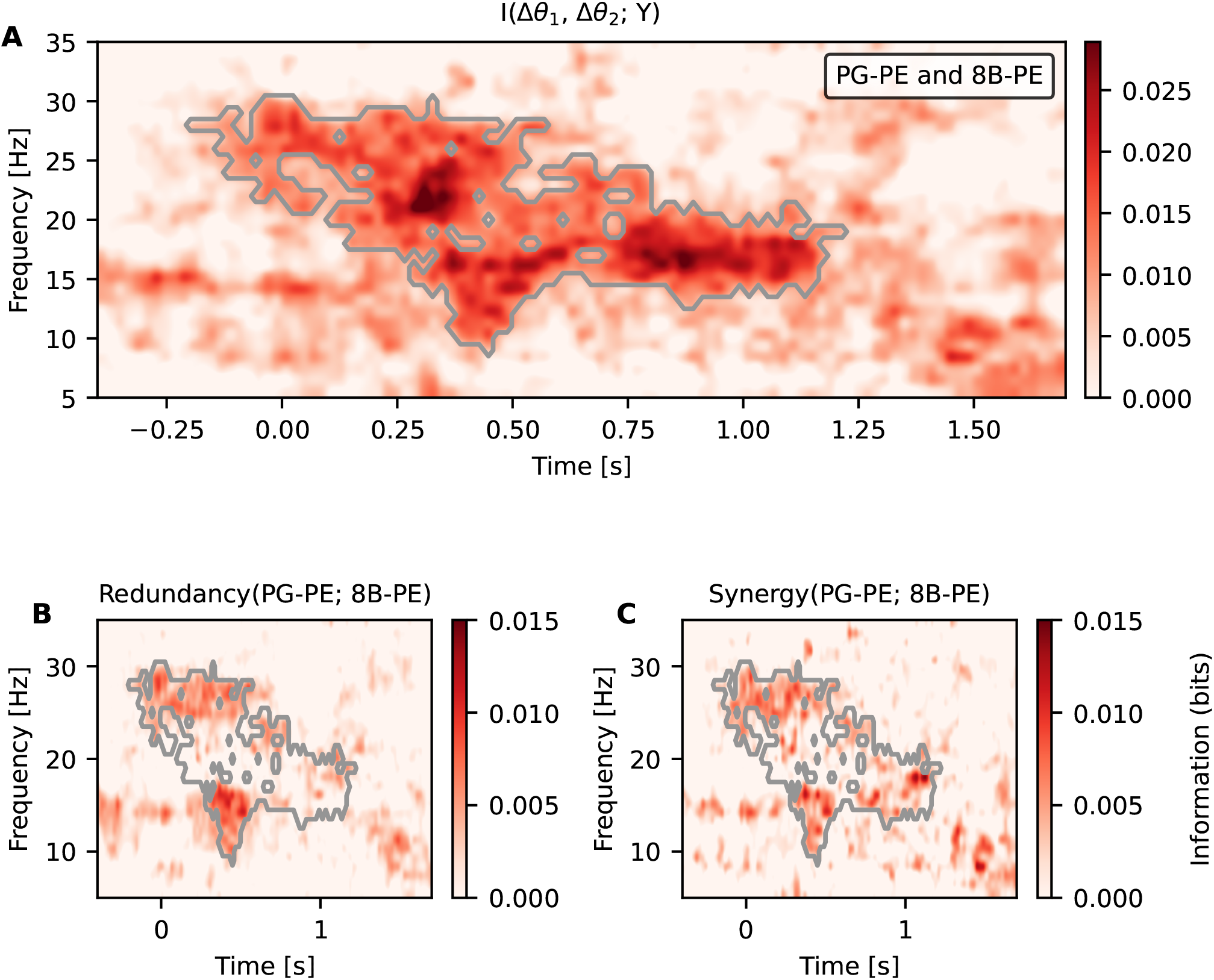
High-order phase relations and amplitude co-modulations during working memory. (**A**) Stimulus-related coherence given by the total mutual-information between phase relations and amplitude co-modulations for parietal-parietal channel pair PG-PE and fronto-parietal channel pair 8B-PE about the stimulus (Eq. 2). The amplitude co-modulation-phase edge for each channel pair is given by Eqs. 9 and 10. (**B**) High-order redundant information about the stimulus contained in the joint amplitude co-modulation and phase-difference components of the signal. Areas delimited by the white line correspond to significant effects obtained via shuffling surrogates (*n* = 1000; *p* < 0.05; corrected using cluster statistics). (**C**) The same as (**B**) but showing the high-order synergistic information.

## DISCUSSION

A large body of evidence supports the view that cognitive functions emerge from the coordinated activity of neural populations distributed across large-scale brain networks (Varela et al., 2001a; Bressler and Menon, 2010), as well as from network-wide, self-organized patterns of information routing (Deco et al., 2015; Battaglia and Brovelli, 2020). However, key questions concerning the underlying neurophysiological mechanisms and computational objectives remain unresolved. For example, how do brain interactions regulate segregation and integration processes to support cognitive functions (Sporns, 2013; Braun, 2015; Deco et al., 2015; Cohen and D’Esposito, 2016; Shine, 2016; Finc, 2020; Wang et al., 2021)? How do functional interactions based on oscillatory synchronization (Engel et al., 2001; Buzsáki and Draguhn, 2004; Fries, 2015) or resonance phenomena (Vinck et al., 2023) encode information about cognitive variables? Do aperiodic transient dynamics and oscillatory phase relations constitute complementary substrates for inter-areal communication and information encoding? How does the brain trades off redundant and synergistic encoding? How do higher-order oscillatory interactions allow distributed encoding over large-scale brain circuits? Progress towards an integrated knowledge in this domain has been limited by the lack of computational tools. In the current study, we provide a novel approach that allows the quantification and decomposition of information about cognitive variables in neural oscillatory networks.

### Neural Oscillatory Partial Information Decomposition (NeOPID)

The first contribution of the study consists in having developed a coherent approach for the decomposition of information about task variables contained in interacting oscillatory time series combining information-theoretical and signal processing approaches. In other words, the proposed approach, termed Neural Oscillatory Partial Information Decomposition (NeOPID), quantifies task-related information in the frequency domain contained in oscillatory interactions. NeOPID integrates Partial Information Decomposition (PID) framework (Williams and Beer, 2010; Wibral et al., 2017) and spectral analysis. This integration yields a set of formal rules for decomposing multivariate mutual information between a set of predictors (i.e., brain signals) and a target variable (e.g., a task variable), and defines unique, redundant, and synergistic information components for oscillatory signals. Although, the formalism is general and may be applicable to other domains involving oscillatory complex systems, the present approach focuses on neural oscillations. Therefore, several assumptions could be made to simplify the inference of information theoretical metrics. First, we adopted the *minimum mutual information* (MMI) PID, which provides accurate estimates for a broad class of systems consistent with a multivariate Gaussian distribution (Barrett, 2015). This allows the inference of redundancy and synergistic terms of PID efficient even for large network. In addition, mutual information and entropy were estimated using Gaussian Copula Mutual Information (GCMI) (Ince, 2017), a semi-parametric, binning-free method that is computationally efficient for the short time series typically encountered in neurophysiological data.

### Phase relations and amplitude co-modulations as complementary and interrelated coding schemes

We first validated our approach through simulations based on the Kuramoto models of coupled oscillators, which is a classical framework for studying oscillatory networks. The analysis of the Kuramoto-based simulations showed that NeOPID can capture the amount of information contained in phase relations between nodes about a task-related variable (Fig. 1). The time-frequency representation of the mutual information between the phase difference of two nodes (Fig. 1F) provides an additional explanatory dimension with respect to the standard spectral coherence (Fig. 1E), which primarily reflects task-unrelated synchronization. The comparison between spectral mutual information and coherence therefore provides a way for assessing the functional role of phase synchronization in relation to cognitive demands and task manipulations.

We then tested the NeOPID approach on simulations based on Stuart–Landau models of coupled oscillators, which extends the Kuramoto model by incorporating both phase and amplitude, thus providing a minimal system capable of exhibiting subcritical Hopf bifurcations, as well as transient and steady-state oscillations (Ponce-Alvarez and Deco, 2024). In the simulations based on a pairwise SL model (Fig. 3A), we observed that amplitude co-modulation contains the dominant fraction of task-related information (Fig. 3C and E). However, we also observed that phase and amplitude co-modulation can interact both redundantly and synergistically (Fig. 3D), suggesting that the degree of interdependence between the two coding schemes can be quantified. Crucially, the NeOPID analysis revealed a synergistic interaction between amplitude co-modulation and phase (Fig. 3D), suggesting that task-related information is not merely present in parallel in these two channels, but it can partly emerge from their joint co-variation in ways not predictable from either alone. In more biologically realistic Stuart-Landau model with multiple oscillators coupled according to macaque monkey structural connectivity (Markov et al., 2014), we observed that inter-areal interactions through synchronization and amplitude co-modulations can track complementary unique information (Fig. 4C and E). For some pairs of areas, the coupling modulation was primarily driven by amplitude co-modulation, with phase relations providing no unique informative contribution (Fig. 4F and H). However, we also observed that phase relations and amplitude co-modulations contain redundant and synergistic information about task variables (Fig. 4D), or be independent (Fig. 4G).

The whole-brain Stuart–Landau network model (Fig. 4) suggests a distance-dependent pattern: proximal, strongly connected areas (V1–V2) tend to show mixed phase–amplitude information, whereas distant regions (V1–area 24c) showed predominantly amplitude-based information with negligible phase contribution. This may suggest the presence of a gradient constrained by long-range projections, which may reduce phase relations due to long conduction delays and thus limiting precise phase locking across distant regions (Buzsáki and Draguhn, 2004; Bastos et al., 2015). Under these conditions, amplitude envelope correlations, being more robust to latency variability, may support long-range information transfer, as observed in large-scale MEG and SEEG studies (Combrisson et al., 2025; Brovelli et al., 2015; Combrisson et al., 2024)). Nevertheless, higher-order analyses (Figs. 5) showed that long-range interactions based on phase-relations can be observed in the whole-brain model, and indicating that distributed cortical representations cannot be reduced to amplitude co-modulations only.

The presence of synergistic information implies that treating phase synchrony and amplitude comodulations as independent proxies for neural coordination (Lachaux et al., 1999; Siegel et al., 2012) may underestimate the information available in their interdependence. More broadly, this finding is consistent with recent theoretical proposals that aperiodic transient dynamics and oscillatory phase relations constitute complementary substrates of inter-areal communication (Vinck et al., 2023).

Taken together, a central finding across our simulations was that amplitude co-modulation and phase relations can provide complementary encoding schemes of task variables, whose contributions can be disentangled and quantified using our approach. The quantification of the degree of redundant, synergistic and unique information present in each coding scheme, may be used to test hypotheses about the role of inter-areal interactions via either oscillatory synchronization (Engel et al., 2001; Buzsáki and Draguhn, 2004; Fries, 2015) or transients resonance phenomena (Vinck et al., 2025) in cognition. In addition, our approach may profit theoretical development of the Communication Through Coherence (CTC) hypothesis (Fries, 2005, 2015), which posits that phase synchrony between neural populations is the primary mechanism for routing and encoding cognitive information. Future analyses using our approach may provide a nuanced perspective on this claim.

### Distributed and higher-order information about task variables via redundant and synergistic oscillatory interactions

A notable contribution of NeOPID is the possibility to investigate the role of distributed and higher-order oscillatory relations via redundant and synergistic encoding in relation to the underlying structural connectivity. We first simulated modulation of neuronal interactions through changes in neuronal synchronization and coupling strength in a network with three nodes using Kuramoto coupled oscillators (Fig. 2). We used different topological organizations of an oscillatory circuit to determine whether information is reflected redundantly or synergistically across interacting edges. A feedforward chain architecture produces predominantly redundant encoding (Fig. 2A), while a convergent collider structure produces synergistic encoding (Fig. 2C). These simulations may provide a direct link between graph-theoretic descriptions of connectivity motifs and information-theoretic characterizations of neural coding. A predominant redundant encoding, as produced by chain architectures resembling feedforward sensory hierarchies, may confer robustness against signal degradation. This may be identified as a predominant redundant information contained in pairs of edges interacting via phase relations, which would reflect higher-order interactions among the phase-difference edges spanning three areas (Fig. 2B). The computational advantage of such coding scheme would be that information about cognitive variable propagates across multiple downstream pathways, so to ensures that no single pathway failure eliminates the representation. In addition, our simulations showed that synergistic encoding, however, as produced by convergence topology (Fig. 2C), was also detectable as a predominant synergistic information in the time-frequency representation (Fig. 2D). A predominant synergistic encoding, as produced by converging architectures resembling top-down influence, may allow functional integration and flexibility properties to the network (Voges et al., 2024). Nevertheless, in anatomically realistic three-node circuits modeled on the V1-V2-V4 circuit, we observed a balanced mixture of both encoding strategies (Fig. 2E-F). In more biologically realistic Stuart-Landau model with multiple oscillators coupled according to macaque monkey structural connectivity, we observed that oscillatory HOI between pairs of edges display both redundant and synergistic information about task modulation via phase relations (Fig. 5), which can be interpreted as a correlate of distributed information about cognitive variables via collective phase relations.

Overall, these results provide evidence that distributed information about cognitive variables via phase relations between multiple brain areas can be extracted from oscillatory activity, and exploit a balance between redundant and synergistic coding schemes. Indeed, the balance between these coding schemes may underpin large-scale robustness and resilience in oscillatory brain network. Synergistic interactions may confer functional advantages over redundancy and unique information by enabling combinatorial coding across brain regions, potentially scaling exponentially with system size Rosas et al. (2020). In contrast, redundancy may support robustness by ensuring that information remains available despite perturbations, at the cost of over-representation, whereas synergy is more vulnerable to disruptions affecting constituent nodes (Mediano, 2022). This would be consistent with the functional advantages of combining robustness and flexibility in distributed cortical circuits (Luppi et al., 2022; Combrisson et al., 2024). More generally, we suggest that the study of the trade-off between redundant and synergistic information about cognitive variables in oscillatory networks may provide a better understanding of the mechanisms regulating segregation and integration in cognition (Sporns, 2013; Braun, 2015; Deco et al., 2015; Cohen and D’Esposito, 2016; Shine, 2016; Finc, 2020; Wang, 2021). To conclude, our study showed that the NeOPID approach can be used as a tool not only for detecting the presence of higher-order oscillatory interactions, but additionally for disentangling their computational function in terms of collective encoding strategies.

### Proof-of-concept on LFPs from fronto-parietal network during working memory

Finally, we provided a proof-of-concept on local field potentials (LFPs) recorded from macaque parietal and premotor cortical regions during a working memory task. We searched for information contained in beta oscillations (15-30Hz) about the stimulus identity during delayed-matched-to-sample task. For an exemplar session, we observed a stimulus-dependent modulation of total mutual information after stimulus onset and during the delay period of the task (Fig. 6A), thus confirming previous findings showing content-specific beta-band fronto-parietal synchronization during visual working memory (Salazar et al., 2012). The total mutual information was decomposed into unique contributions due to amplitude co-modulation and phase relations. For the session under investigation, the results show that most of the information about stimulus was contained in amplitude co-modulations (Fig. 6B), rather than phase relations (Fig. 6C). This suggests that future studies may profit from disentangling synchrony- and amplitude-based information components.

The higher-order analysis (Fig. 7) further revealed that pairs of fronto-parietal recording sites jointly reflect stimulus information through both redundant and synergistic interactions during the memory delay period. The synergistic component was particularly noteworthy: it implies that certain aspects of the memorized stimulus are only jointly contained by multiple fronto-parietal channel pairs simultaneously. However, the result confirmed the prediction from whole-brain simulations, suggesting a balanced encoding based on redundancy and synergistic information. The co-existence of redundant and synergistic higher-order encoding within the same fronto-parietal network suggests that the same oscillatory system simultaneously supports reliable broadcast and flexible integration of working memory content, an organizational principle with direct implications for biologically plausible models of working memory maintenance (Dotson et al., 2018; Miller et al., 2018).

### Methodological considerations

Recent advances in information theory have provided tools that share important conceptual foundations with our implementation of NeOPID. Faes et al. (2021) introduced a spectral decomposition of multivariate information measures for physiological oscillations, yielding frequency-specific quantification of unique, redundant, and synergistic information between cardiovascular and cardiorespiratory signals, the same partial information decomposition (PID) formalism adopted by NeOPID. Building on this, Faes et al. (2022) and Sparacino et al. (2023) extended frequency-domain information decomposition to higher-order interactions (HOIs) beyond pairwise relations, introducing the O-information rate (OIR) to characterize the balance between redundancy and synergy among groups of three or more oscillatory processes in both the time and frequency domains. Most recently, Antonacci et al. (2025) further generalized this framework to non-stationary systems by combining time-varying vector autoregressive models with recursive estimation, enabling time-frequency tracking of transient HOIs in neural recordings. The central novelty of the use of NeOPID with respect to these prior tools lies in the nature of the variable being decoded. Rather than quantifying the nature of oscillatory interactions, NeOPID quantifies the information about external task or cognitive variables contained in the interactions between signals. This distinction maps onto the difference between functional connectivity (signal-to-signal statistical relations) and information encoding (brain-behavior dependence). An important direction for future work is to integrate these two perspectives, combining the study of multivariate information in the frequency domain in complex oscillatory systems with the task-variable decomposition of NeOPID, so to obtain a unified account of how intrinsic oscillatory interactions are reflected in external task variables. An additional potential extension is the study of across-orders interactions from pairwise to higher-order interactions in oscillatory networks, with the potential application to multiscale systems.

### Potential limitations

Several aspects of the current implementation merit consideration. First, the minimum mutual information (MMI) PID estimator (Barrett, 2015), while computationally tractable and well-calibrated for systems with multivariate Gaussian statistics, provides only a lower bound on synergistic information and may not faithfully decompose the unique information atoms across all probability regimes (Williams and Beer, 2010; Wibral et al., 2017). Future implementations could benefit from more principled lattice-based PID comparing different approaches for information decomposing (Liardi et al., 2026). Second, the Gaussian-copula mutual information estimator (Ince, 2017) provides a robust and sample-efficient approximation for short neural data series, but its reliance on rank-based normalization to Gaussian may underestimate information contained in non-monotonic relationships between signal features and task variables. Finally, the neurophysiological results reported here are based on a single session from a single animal, and therefore constitute a proof-of-concept; validation across multiple animals, sessions, and task designs is necessary to establish the generality of encoding principles.

## Future directions

Several extensions of NeOPID follow directly from these results. First, the current formulation is symmetric and does not estimate directionality; incorporating measures grounded in the Wiener–Granger causality principle, such as the Feature-specific Information Transfer (Celotto, 2024) would allow detection of asymmetric interactions between oscillatory signals. Second, extending the framework to cross-frequency interactions (e.g., theta phase–gamma amplitude coupling in hippocampo-cortical circuits) (Jensen and Colgin, 2007; Canolty and Knight, 2010) would enable testing whether multiplexed frequency codes carry unique or synergistic task information and clarify the computational role of cross-frequency nesting. Finally, the framework could be used to study whether differential roles of broadband and oscillatory dynamics in encoding visual information (Roberts et al., 2026) is carried by phase alignment and/or amplitude co-modulation, or their interaction.

## Conclusions

The Neural Oscillatory Partial Information Decomposition (NeOPID) framework provides tools to characterize how oscillatory neural interactions encode cognitive information. It quantifies complementary and synergistic contributions of phase synchrony and amplitude co-modulation, and extends redundancy/synergy analyses to higher-order interactions, thereby expanding methods for studying distributed oscillatory coding. Validation across canonical models, whole-brain simulations, and beta-band activity in fronto-parietal networks supports its use for probing how cognitive variables are represented in distributed oscillatory systems.

## Acknowledgments

A.B. and V.L. were supported by the A*Midex Foundation of Aix-Marseille University project “Hinteract” (AMX-22-RE-AB-071). A.B, D.M. were supported by EU’s Horizon 2020 Framework Programme for Research and Innovation under the Specific Grant Agreements No. 101147319 (EBRAINS 2.0 Project). The “Center de Calcul Intensif of the Aix-Marseille University (CCIAM)” is acknowledged for high-performance computing resources. We are grateful to Dr. Charles M. Gray (Montana University) for his helpful comments on earlier versions of the manuscript.

## References

Antonacci Y, Bara’ C, Sparacino L, Mijatovic G, Minati L, Faes L (2025) A method for the time-frequency analysis of high-order interactions in non-stationary physiological networks.

Barrett AB (2015) Exploration of synergistic and redundant information sharing in static and dynamical gaussian systems. Phys. Rev. E Stat. Nonlin. Soft Matter Phys. 91.

Bastos AM, Vezoli J, Bosman CA, Schoffelen JM, Oostenveld R, Dowdall JR, De Weerd P, Kennedy H, Fries P (2015) Visual areas exert feedforward and feedback influences through distinct frequency channels. Neuron 85:390–401.

Battaglia D, Brovelli A (2020) Functional connectivity and neuronal dynamics: insights from computational methods.

Battiston F (2021) The physics of higher-order interactions in complex systems. Nat. Phys. 17.

Battiston F, Cencetti G, Iacopini I, Latora V, Lucas M, Patania A, Young JG, Petri G (2020) Networks beyond pairwise interactions: Structure and dynamics. Physics Reports 874:1–92.

Benson AR, Gleich DF, Leskovec J (2016) Higher-order organization of complex networks. Science 353.

Bonnefond M, Kastner S, Jensen O (2017) Communication between brain areas based on nested oscillations. eneuro 4.

Bosman CA, Schoffelen JM, Brunet N, Oostenveld R, Bastos AM, Womelsdorf T, Rubehn B, Stieglitz T, De Weerd P, Fries P (2012) Attentional stimulus selection through selective synchronization between monkey visual areas. Neuron 75:875–888.

Braun U (2015) Dynamic reconfiguration of frontal brain networks during executive cognition in humans.Proc. Natl. Acad. Sci. USA. 112.

Bressler SL, Menon V (2010) Large-scale brain networks in cognition: emerging methods and principles.Trends Cogn. Sci. 14.

Brovelli A, Chicharro D, Badier JM, Wang H, Jirsa V (2015) Characterization of cortical networks and corticocortical functional connectivity mediating arbitrary visuomotor mapping. J. Neurosci. 35.

Buschman TJ, Miller EK (2007) Top-down versus bottom-up control of attention in the prefrontal and posterior parietal cortices. science 315:1860–1862.

Buzsáki G, Draguhn A (2004) Neuronal oscillations in cortical networks. Science 304.

Canolty RT, Knight RT (2010) The functional role of cross-frequency coupling. Trends Cogn. Sci. 14:506–515.

Celotto M (2024) An information-theoretic quantification of the content of communication between brain regions. Adv. Neural Inf. Process. Syst. 36.

Chelaru MI (2021) High-order interactions explain the collective behavior of cortical populations in executive but not sensory areas. Neuron 109.

Cohen JR, D’Esposito M (2016) The segregation and integration of distinct brain networks and their relationship to cognition. J. Neurosci. 36.

Cohen JR, D’Esposito M (2016) The segregation and integration of distinct brain networks and their relationship to cognition. Journal of Neuroscience 36:12083–12094.

Combrisson E (2022) Group-level inference of information-based measures for the analyses of cognitive brain networks from neurophysiological data. Neuroimage 258.

Combrisson E, Basanisi R, Cordeiro VL, Ince RA, Brovelli A (2022) Frites: A python package for functional connectivity analysis and group-level statistics of neurophysiological data. Journal of Open Source Software 7:3842.

Combrisson E, Basanisi R, Gueguen MCM, Rheims S, Kahane P, Bastin J, Brovelli A (2024) Neural interactions in the human frontal cortex dissociate reward and punishment learning .

Combrisson E, Basanisi R, Neri M, Auzias G, Petri G, Marinazzo D, Panzeri S, Brovelli A (2025) Higher-order and distributed synergistic functional interactions encode information gain in goal-directed learning. Nature Communications 16:7179.

Crutchfield JP (1994) The calculi of emergence: computation, dynamics and induction. Phys. D 75.

Deco G, Tononi G, Boly M, Kringelbach ML (2015) Rethinking segregation and integration: contributions of whole-brain modelling. Nat. Rev. Neurosci. 16.

Dotson NM, Hoffman SJ, Goodell B, Gray CM (2018) Feature-based visual short-term memory is widely distributed and hierarchically organized. Neuron 99:215–226.

Engel AK, Fries P, Singer W (2001) Dynamic predictions: Oscillations and synchrony in top–down processing. Nat. Rev. Neurosci. 2.

Engel AK, Gerloff C, Hilgetag CC, Nolte G (2013) Intrinsic coupling modes: multiscale interactions in ongoing brain activity. Neuron 80:867–886.

Faes L, Mijatovic G, Antonacci Y, Pernice R, Barać C, Sparacino L, Sammartino M, Porta A, Marinazzo D, Stramaglia S (2022) A new framework for the time- and frequency-domain assessment of high-order interactions in networks of random processes. IEEE Transactions on Signal Processing 70:5766–5777.

Faes L, Pernice R, Mijatovic G, Antonacci Y, Krohova JC, Javorka M, Porta A (2021) Information decomposition in the frequency domain: a new framework to study cardiovascular and cardiorespiratory oscillations. Philosophical Transactions of the Royal Society A: Mathematical, Physical and Engineering Sciences 379:20200250.

Finc K (2020) Dynamic reconfiguration of functional brain networks during working memory training. Nat. Commun. 11.

Fisher NI (1995) Statistical analysis of circular data cambridge university press.

Fries P (2015) Rhythms for cognition: communication through coherence. Neuron 88.

Fries P (2005) A mechanism for cognitive dynamics: neuronal communication through neuronal coherence. Trends in cognitive sciences 9:474–480.

Fries P (2009) Neuronal gamma-band synchronization as a fundamental process in cortical computation. Annual review of neuroscience 32:209–224.

Fries P (2015) Rhythms for cognition: communication through coherence. Neuron 88:220–235.

Fries P, Reynolds JH, Rorie AE, Desimone R (2001) Modulation of oscillatory neuronal synchronization by selective visual attention. Science 291:1560–1563.

Grilli J, Barabás G, Michalska-Smith MJ, Allesina S (2017) Higher-order interactions stabilize dynamics in competitive network models. Nature 548.

Ince RAA (2017) A statistical framework for neuroimaging data analysis based on mutual information estimated via a gaussian copula. Hum. Brain Mapp. 38.

Jensen O, Colgin LL (2007) Cross-frequency coupling between neuronal oscillations.

Joglekar MR, Mejias JF, Yang GR, Wang XJ (2018) Inter-areal balanced amplification enhances signal propagation in a large-scale circuit model of the primate cortex. Neuron 98:222–234.

Köster U, Sohl-Dickstein J, Gray CM, Olshausen BA (2014) Modeling higher-order correlations within cortical microcolumns. PLoS Comput Biol. 10.

Lachaux JP, Rodriguez E, Martinerie J, Varela FJ (1999) Measuring phase synchrony in brain signals. Human brain mapping 8:194–208.

Levine JM, Bascompte J, Adler PB, Allesina S (2017) Beyond pairwise mechanisms of species coexistence in complex communities. Nature 546.

Liardi A, Down KJA, Blackburne G, Neri M, Mediano PAM (2026) The mathematical landscape of partial information decomposition: A comprehensive review of properties and measures.

Lizier JT, Flecker B, Williams PL (2013) Towards a synergy-based approach to measuring information modification In 2013 IEEE Symposium on Artificial Life (ALIFE), pp. 43–51. IEEE.

Lizier JT, Prokopenko M, Zomaya AY (2007) Information transfer by particles in cellular automata In Australian Conference on Artificial Life, pp. 49–60. Springer.

Lundqvist M, Rose J, Herman P, Brincat SL, Buschman TJ, Miller EK (2016) Gamma and beta bursts underlie working memory. Neuron 90:152–164.

Luppi AI, Rosas FE, Mediano PAM, Menon DK, Stamatakis EA (2024) Information decomposition and the informational architecture of the brain. Trends Cogn. Sci. 28.

Luppi AI, Mediano PA, Rosas FE, Holland N, Fryer TD, O’Brien JT, Rowe JB, Menon DK, Bor D, Stamatakis EA (2022) A synergistic core for human brain evolution and cognition. Nature Neuroscience 25:771–782.

Markov NT, Ercsey-Ravasz M, Van Essen DC, Knoblauch K, Toroczkai Z, Kennedy H (2013) Cortical high-density counterstream architectures. Science 342:1238406.

Markov NT, Ercsey-Ravasz MM, Ribeiro Gomes A, Lamy C, Magrou L, Vezoli J, Misery P, Falchier A, Quilodran R, Gariel MA et al. (2014) A weighted and directed interareal connectivity matrix for macaque cerebral cortex. Cerebral cortex 24:17–36.

Martignon L (2000) Neural coding: higher-order temporal patterns in the neurostatistics of cell assemblies. Neural Comput 12.

Mediano PAM (2022) Integrated information as a common signature of dynamical and information-processing complexity. Chaos 32.

Miller EK, Lundqvist M, Bastos AM (2018) Working memory 2.0. Neuron 100:463–475.

Neri M, Vinchhi D, Ferreyra C, Robiglio T, Ates O, Ontivero-Ortega M, Brovelli A, Marinazzo D, Combrisson E (2024) Hoi: A python toolbox for high-performance estimation of higher-order interactions from multivariate data. Journal of Open Source Software 9:7360.

Nigam S, Pojoga S, Dragoi V (2019) Synergistic coding of visual information in columnar networks. Neuron 104.

Panzeri S, Moroni M, Safaai H, Harvey CD (2022) The structures and functions of correlations in neural population codes. Nat. Rev. Neurosci. 23.

Ponce-Alvarez A, Deco G (2024) The hopf whole-brain model and its linear approximation. Scientific reports 14:2615.

Roberts L,Äijälä J, Burger F, Uran C, Jensen MA, Miller KJ, Ince RAA, Vinck M, Hermes D, Canales-Johnson A (2026) Broadband synergy versus oscillatory redundancy in the visual cortex. Nat. Commun. 17.

Rosas FE, Mediano PAM, Rassouli B, Barrett AB (2020) An operational information decomposition via synergistic disclosure. J. Phys. A: Math. Theor. 53.

Salazar R, Dotson N, Bressler S, Gray C (2012) Content-specific fronto-parietal synchronization during visual working memory. Science 338:1097–1100.

Sanchez-Gorostiaga A, Bajić D, Osborne ML, Poyatos JF, Sanchez A (2019) High-order interactions distort the functional landscape of microbial consortia. PLoS Biol. 17.

Santoro A, Battiston F, Lucas M, Petri G, Amico E (2024) Higher-order connectomics of human brain function reveals local topological signatures of task decoding, individual identification, and behavior. Nat. Commun. 15.

Santoro A, Battiston F, Petri G, Amico E (2023) Higher-order organization of multivariate time series. Nature Physics 19:221–229.

Schneidman E, Berry MJ, Segev R, Bialek W (2006) Weak pairwise correlations imply strongly correlated network states in a neural population. Nature 440.

Shahidi N, Andrei AR, Hu M, Dragoi V (2019) High-order coordination of cortical spiking activity modulates perceptual accuracy. Nat. Neurosci. 22.

Shine JM (2016) The dynamics of functional brain networks: integrated network states during cognitive task performance. Neuron 92.

Siegel M, Donner TH, Engel AK (2012) Spectral fingerprints of large-scale neuronal interactions. Nature Reviews Neuroscience 13:121–134.

Singer W, Gray CM (1995) Visual feature integration and the temporal correlation hypothesis. Annual review of neuroscience 18:555–586.

Sparacino L, Antonacci Y, Marinazzo D, Stramaglia S, Faes L (2023) Quantifying high-order interactions in complex physiological networks: A frequency-specific approach In Cherifi H, Mantegna RN, Rocha LM, Cherifi C, Micciché S, editors, Complex Networks and Their Applications XI, pp. 301–309, Cham. Springer International Publishing.

Sporns O (2013) Network attributes for segregation and integration in the human brain. Curr. Opin. Neurobiol. 23.

Valente M (2021) Correlations enhance the behavioral readout of neural population activity in association cortex. Nat. Neurosci. 24.

van Kerkoerle T, Self MW, Dagnino B, Gariel-Mathis MA, Poort J, van der Togt C, Roelfsema PR (2014) Alpha and gamma oscillations characterize feedback and feedforward processing in monkey visual cortex. Proc. Natl. Acad. Sci. U. S. A. 111:14332–14341.

Varela F, Lachaux JP, Rodriguez E, Martinerie J (2001a) The brainweb: phase synchronization and large-scale integration. Nat. Rev. Neurosci. 2.

Varela F, Lachaux JP, Rodriguez E, Martinerie J (2001b) The brainweb: phase synchronization and large-scale integration. Nature reviews neuroscience 2:229–239.

Varley TF, Pope M, Faskowitz J, Sporns O (2023a) Multivariate information theory uncovers synergistic subsystems of the human cerebral cortex. Communications biology 6:451.

Varley TF, Pope M, Grazia M, Joshua Sporns O (2023b) Partial entropy decomposition reveals higher-order information structures in human brain activity. Proceedings of the National Academy of Sciences 120:e2300888120.

Vezoli J, Vinck M, Bosman CA, Bastos AM, Lewis CM, Kennedy H, Fries P (2021) Brain rhythms define distinct interaction networks with differential dependence on anatomy. Neuron 109:3862–3878.

Vinck M, Uran C, Dowdall JR, Rummell B, Canales-Johnson A (2025) Large-scale interactions in predictive processing: oscillatory versus transient dynamics. Trends Cogn. Sci. 29.

Vinck M, Uran C, Spyropoulos G, Onorato I, Broggini AC, Schneider M, Canales-Johnson A (2023) Principles of large-scale neural interactions. Neuron 111:987–1002.

Vinck M, Van Wingerden M, Womelsdorf T, Fries P, Pennartz CM (2010) The pairwise phase consistency: a bias-free measure of rhythmic neuronal synchronization. Neuroimage 51:112–122.

Voges N, Lima V, Hausmann J, Brovelli A, Battaglia D (2024) Decomposing neural circuit function into information processing primitives. Journal of Neuroscience 44.

Wang R (2021) Segregation, integration, and balance of large-scale resting brain networks configure different cognitive abilities. Proc. Natl. Acad. Sci. USA. 118.

Wang R, Liu M, Cheng X, Wu Y, Hildebrandt A, Zhou C (2021) Segregation, integration, and balance of large-scale resting brain networks configure different cognitive abilities. Proceedings of the National Academy of Sciences 118:e2022288118.

Wibral M, Priesemann V, Kay JW, Lizier JT, Phillips WA (2017) Partial information decomposition as a unified approach to the specification of neural goal functions. Brain Cogn. 112.

Williams PL, Beer RD (2010) Nonnegative decomposition of multivariate information. arXiv .

Womelsdorf T, Fries P (2007) The role of neuronal synchronization in selective attention. Current opinion in neurobiology 17:154–160.

Womelsdorf T, Schoffelen JM, Oostenveld R, Singer W, Desimone R, Engel AK, Fries P (2007) Modulation of neuronal interactions through neuronal synchronization. science 316:1609–1612.

Xiong YS, Donoghue JA, Lundqvist M, Mahnke M, Major AJ, Brown EN, Miller EK, Bastos AM (2024) Propofol-mediated loss of consciousness disrupts predictive routing and local field phase modulation of neural activity. Proceedings of the National Academy of Sciences 121:e2315160121.

Yu S (2011) Higher-order interactions characterized in cortical activity. J. Neurosci. 31.

